# Identification of Iron-Sulfur (Fe-S) and Zn-binding Sites Within Proteomes Predicted by DeepMind’s AlphaFold2 Program Dramatically Expands the Metalloproteome

**DOI:** 10.1101/2021.10.08.463726

**Authors:** Zachary J. Wehrspan, Robert T. McDonnell, Adrian H. Elcock

**Affiliations:** Department of Biochemistry, University of Iowa, Iowa City, Iowa, USA

**Keywords:** Ligands, metalloproteomics, functional annotation, UniProt

## Abstract

DeepMind’s AlphaFold2 software has ushered in a revolution in high quality, 3D protein structure prediction. In very recent work by the DeepMind team, structure predictions have been made for entire proteomes of twenty-one organisms, with >360,000 structures made available for download. Here we show that thousands of novel binding sites for iron-sulfur (Fe-S) clusters and zinc ions can be identified within these predicted structures by exhaustive enumeration of all potential ligand-binding orientations. We demonstrate that AlphaFold2 routinely makes highly specific predictions of ligand binding sites: for example, binding sites that are comprised exclusively of four cysteine sidechains fall into three clusters, representing binding sites for 4Fe-4S clusters, 2Fe-2S clusters, or individual Zn ions. We show further: (a) that the majority of known Fe-S cluster and Zn-binding sites documented in UniProt are recovered by the AlphaFold2 structures, (b) that there are occasional disputes between AlphaFold2 and UniProt with AlphaFold2 predicting highly plausible alternative binding sites, (c) that the Fe-S cluster binding sites that we identify in *E. coli* agree well with previous bioinformatics predictions, (d) that cysteines predicted here to be part of Fe-S cluster or Zn-binding sites show little overlap with those shown via chemoproteomics techniques to be highly reactive, and (e) that AlphaFold2 occasionally appears to build erroneous disulfide bonds between cysteines that should instead coordinate a ligand. These results suggest that AlphaFold2 could be an important tool for the functional annotation of proteomes, and the methodology presented here is likely to be useful for predicting other ligand-binding sites.

## Introduction

Approximately twenty-five years ago, structural genomics initiatives dramatically increased the known universe of protein structures [1]. The rapid deposition of large numbers of experimentally determined protein structures in the Protein Data Bank [2], many of which were of unknown function, drove the development of a variety of computational methods for predicting functionally important regions within proteins from structure alone (e.g. [3–5]); interesting developments continue to this day (e.g. [6]). In the intervening years, the accuracy of protein structure prediction using computational methods has steadily improved [7], with a sudden, extraordinary increase in accuracy achieved at the most recent CASP14 meeting by DeepMind’s AlphaFold2 method [8]. Very recently, the DeepMind team described their method in detail [8] and demonstrated its potential utility by reporting structures for almost complete proteomes of 21 organisms [9]. The unprecedented sudden availability of hundreds of thousands of new (predicted) protein structures provides a host of new opportunities for the functional annotation of proteomes.

One functional attribute of a protein that is important to determine is whether it binds cofactors or other ligands, so an immediate question to ask is the extent to which analysis of the AlphaFold2 structures might allow recognition of potential ligand binding sites. Interestingly, the DeepMind team has already provided one clear example of a protein whose predicted structure contains a known zinc (Zn) binding site that their AlphaFold2 method has been able to automatically construct [8]. Most impressively, the binding site had the four coordinating cysteine sidechains oriented tetrahedrally to provide a near-ideal binding site for the metal, even though the structure prediction process was conducted in the absence of the metal ion. This behavior reflects the fact that AlphaFold2’s side chain predictions have been shown to be extremely accurate if the underlying backbone conformation is correct [8].

Here, we aim to expand on that intriguing result with a focus on identifying potential binding sites for Fe-S clusters and Zn ions within the full set of 362,311 protein structures made available by the DeepMind team. We have selected Fe-S clusters and Zn ions as ligands for the following reasons. First, both are biologically significant: Fe-S clusters play important biological roles in electron transfer, catalysis, and sensing iron and oxygen [10–12], while Zn binding is important for stabilizing structurally diverse zinc-finger domains in proteins involved in an extraordinarily wide variety of cellular processes [13, 14].

Second, a successful computational method for identifying binding sites for Fe-S clusters and Zn ions in proteins would be a valuable complement to more involved and expensive experimental techniques that seek to identify the same. Chemoproteomics experiments, for example, have allowed Fe-S cluster binding proteins in *E. coli* [15] and Zn binding proteins in human cancer cells [16] to be identified on a proteomic level; such experiments exploit the differential chemical reactivity of coordinating cysteines under conditions in which the ligand is abundant or scarce. Two practical disadvantages with such experiments are: (a) that proteins whose expression levels are low under the studied growth conditions can evade detection, and (b) that binding sites that are deeply buried within a protein’s structure and therefore difficult for reactive agents to access, might also remain unidentified [15]. One potential major advantage of computational methods, therefore, is that they are unaffected by both issues.

A final reason for choosing Fe-S cluster binding sites and Zn binding sites is that they have well defined binding sites. Both types of binding site have been extensively documented in the literature, with the geometric characteristics and the coordinating residues thoroughly analyzed in a number of structural databases, e.g. the comprehensive MetalPDB database [17] which attempts to document all metal binding sites in macromolecules, and ZincBind, a database specifically focused on Zn binding sites [18], and with webservers dedicated to determining and validating metal coordination geometries in crystal structures (e.g. FindGeo [19] and CheckMyMetal [20]). A number of studies have attempted to exploit structural data, often in combination with bioinformatics methods, in an attempt to identify potential Fe-S cluster-binding proteins on a proteomic scale (e.g. [21, 22]) and related methods have been developed with a view to predicting Zn-binding sites (e.g. [23]) and, very recently, to attempt to predict whether a metal-binding site within a protein is likely to be catalytically active or inactive [24].

Guided by previous studies detailing both the prevalence of different Fe-S cluster types and the different common arrangements of coordinating residues around both Fe-S clusters and isolated Zn ions, we seek here to identify potential binding sites for both types of ligands by exhaustively placing them at all plausible locations within the structures. While similar template-matching approaches have been used previously to find ligand binding sites in crystallographic structures (e.g. [25–27]), the DeepMind team’s dataset now offers the opportunity to extend such searches to 21 near-complete proteomes. Our results indicate that thousands of highly plausible binding sites can be identified for Fe-S clusters and Zn ions, thus expanding significantly the size of the known metalloproteome. If AlphaFold2’s extraordinary ability to automatically generate realistic binding sites extends to other classes of biologically important ligands, AlphaFold2 could become an important tool for functionally annotating proteomes.

## Results

### A ligand-search algorithm can identify Fe-S- and Zn-binding sites in AlphaFold2 structures

Our protocol for identifying potential binding sites for Fe-S clusters and Zn ions is illustrated schematically in Figure 1a. Briefly, we start by compiling a list of “ligand types” whose binding sites we wish to search for; in the present study, six of these ligand types are variants of Fe-S clusters and six are variants of Zn binding sites; see Methods). Our code then identifies all possible “regions” in the protein structure where binding sites for these ligand types might be found, and examines each such region in turn, exhaustively sampling all possible superpositions of each ligand type onto all potential coordinating residues. The largest region such that we found in the human protein contained 107 potential coordinating residues and 194 atoms. Ligands are added to regions, according to their priority in the list of ligand types, if: (a) the root mean square deviation (RMSD) of their superposition is below a desired threshold (i.e. 0.5 Å in most of the cases discussed here), and (b) they are free of steric clashes. Ligands are iteratively added within each region until no further success is achieved, at which point the protocol moves on to the next region, repeating the process until all regions in the protein have been examined. Remarkably, binding sites within the AlphaFold2 structures are, in a large number of cases, sufficiently well-formed that fits of ligands within them result immediately in structures that are highly credible. Figure 1b, for example, shows a representative binding site for a 4Fe-4S cluster that was identified, together with its coordinating cysteines; Figure 1c displays an example in which two nearby Zn binding sites were identified within the same region, each sharing one coordinating cysteine and one coordinating histidine; this particular example echoes a simpler case shown by the DeepMind team in the AlphaFold2 methodological paper [8].

**Figure 1.**
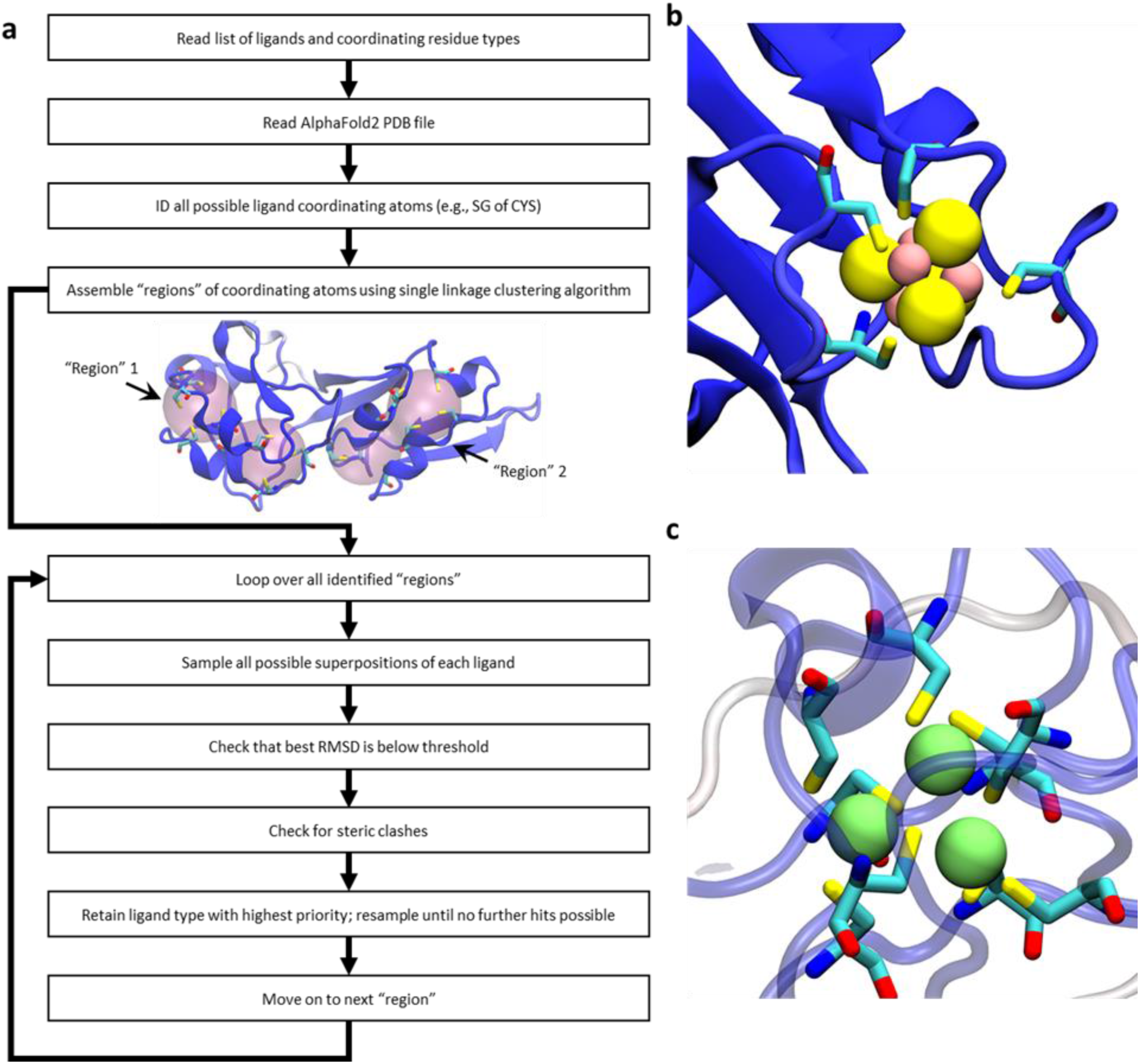
Schematic illustration of our ligand-search protocol and examples of identified metal binding sites in AlphaFold2 structures. (a) a flowchart outlining the method. For the protein illustrated in the center of the flowchart, two “regions” are highlighted; these are distinct from one another, and are treated as separate entities when enumerating all possible ligand superpositions. (b) an example of an identified 4Fe-4S cluster coordinated by four cysteine residues (ligand type “4Fe-4S Cys_4_”; UniProt accession: <u>**P08201**</u>). (c) an example of a cluster of three closely spaced Zn binding sites coordinated by a total of eight cysteine residues; this cluster is formed by three successful additions of the ligand type “Zn Cys_4_” (UniProt accession: <u>**P05100**</u>). In all images of proteins in this figure, the protein is colored by each residue’s pLDDT score with dark blue indicating a “very high confidence” prediction by AlphaFold2 and dark red indicting a “low confidence” prediction.

### Binding sites identified within AlphaFold2 structures can be unambiguously assigned to one ligand type

Since our primary criterion for identifying a potential ligand binding site is the RMSD obtained when the coordinating atoms are superimposed, it is important to determine whether this can unambiguously identify a site as binding one particular ligand in preference to others. The simplest way to illustrate this comes from examining all regions in the entire dataset that contain only four cysteine side chains. We consider such regions as potential binding sites for: (1) a 4Fe-4S cluster, (2) a 2Fe-2S cluster, and (3) a single Zn ion (see Methods). For each such region, we calculate a best-RMSD value for each of the three possible ligand types and we plot all datapoints for which any of these values satisfies our 0.5 Å RMSD threshold in Figure 2a (points are colored according to their “pLDDT score”, which is AlphaFold2’s measure of prediction confidence; see below). Strikingly, three regions of high density are obvious, each corresponding to a well-defined binding site for one of the three ligand types. For example, the group of points at the top-right of Figure 2a represents the 148 cases that are identified as binding sites for 4Fe-4S clusters. The mean RMSD coordinates of these cases are: 0.28, 2.01, 1.44 Å. In structural terms, therefore, these are excellent binding sites for 4Fe-4S clusters (mean RMSD 0.28 Å), but poor binding sites for 2Fe-2S clusters (mean RMSD 2.01 Å), and single Zn ions (mean RMSD 1.44 Å). Similarly, the group of points at the top-left of Figure 2a represent the 127 cases that are identified as binding sites for 2Fe-2S clusters (mean RMSD coordinates of: 2.15, 0.32, 1.78 Å), while the group of points at the bottom of Figure 2a represent the 5344 cases that are predicted to be binding sites for isolated Zn ions (mean RMSD coordinates of: 1.64, 1.82, 0.17 Å). The well-separated nature of each of the three groups indicates that, in all of the cases where AlphaFold2 builds a plausible ligand binding site, it makes a clear decision about which type of binding site to build: it does not simply bring four cysteine sidechains together in a non-specific structural arrangement.

**Figure 2.**
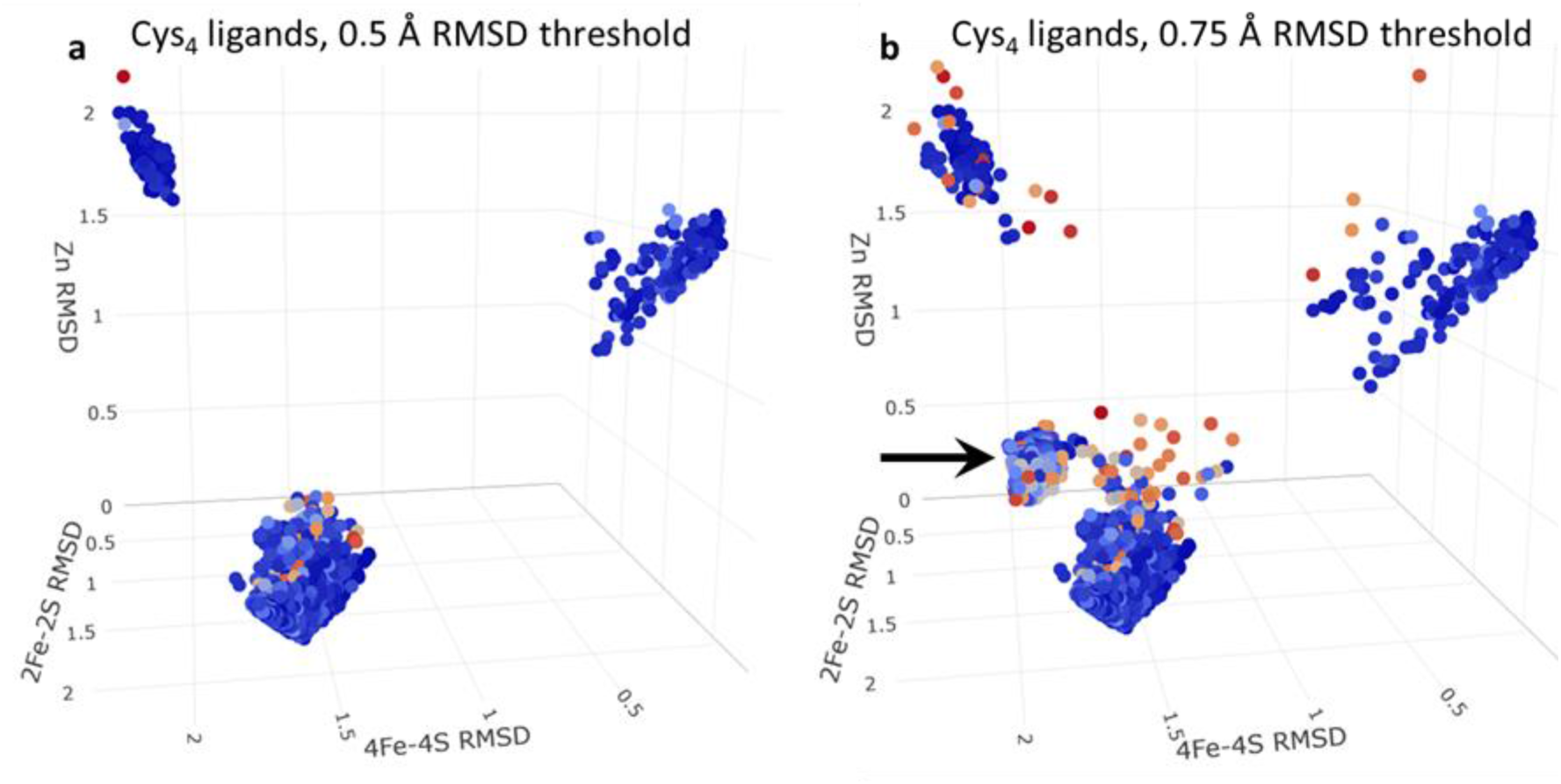
3D scatterplot of the RMSDs of successfully placed ligand types coordinated by 4 cysteine residues. Each point represents a single ligand placed successfully and colored by the minimum pLDDT score of any of its coordinating residues; each axis represents the RMSD of the fit of one of the three possible ligand types (“4Fe-4S Cys_4_”, “2Fe-2S Cys_4_”, “Zn Cys_4_”) on to the coordinating cysteines. (a) shows all ligands that pass an RMSD threshold of 0.5 Å. (b) shows all ligands that pass an RMSD threshold of 0.75 Å. The black arrow highlights the distinct grouping of Zn ligands that appear to contain erroneous disulfide bonds.

Interestingly, if we relax the RMSD threshold for defining potential new binding sites from 0.5 Å to 0.75 Å (Figure 2b), a fourth group of points appears (see black arrow) with mean RMSD coordinates 1.93, 1.67, 0.69 Å). Visual examination of a number of these cases indicates that these tend to be malformed Zn binding sites in which two of the four cysteines have been disulfide-bonded to each other. We suspect that in most cases these disulfide bonds are erroneous, and that such cases are ones where AlphaFold2 is unable to make a clear decision about what to build. Supporting that interpretation, it is notable that the pLDDT scores of the residues constituting such binding sites are somewhat lower (i.e. lower confidence) than those of the three conventional binding sites identified above: the former have a mean minimum pLDDT score of 79 ± 8 (N = 470), while the latter have mean minimum pLDDT scores of 90 ± 9 (N = 180), 91 ± 16 (N = 163), 89 ± 8 (N = 5296) for 4Fe-4S, 2Fe-2S, and Zn binding sites, respectively. Importantly, such cases can be eliminated by restricting attention only to those binding sites that satisfy the more stringent RMSD threshold of 0.5 Å.

The notion that AlphaFold2 tends to build binding sites that are specific for one selected type of ligand is supported by the distribution of RMSD values obtained for all successfully placed instances of each of the twelve ligand types (Figure S1). In most cases, the RMSD values peak at values at or below 0.2 Å, indicating that the identified binding site is essentially a perfect match for that particular ligand; the only cases where substantial populations of less-ideal fits occur (i.e. ones with RMSDs that approach the threshold value of 0.5 Å) are with the deliberately more relaxed ligand types that contain only three points of superposition (see Methods).

### Many AlphaFold2 structures are predicted to contain multiple Fe-S clusters and/or Zn ions

In addition to generating binding sites that are unambiguously intended for specific ligands (e.g. a 4Fe-4S cluster in preference to a 2Fe-2S cluster), AlphaFold2 also routinely builds multiple ligand binding sites within an individual protein. Figure 3a shows a histogram of the number of binding sites identified within each of the 362,311 proteins for which AlphaFold2 predictions have been made available. The frequencies are plotted on a logarithmic scale, with separate histograms shown for those proteins identified as binding Fe-S clusters (in red) and those identified as binding one or more Zn ions (in blue). Both histograms are broad and long-tailed, with the most extreme cases being a protein in *M. jannaschii* (uncharacterized polyferredoxin-like protein MJ1303; UniProt Accession: <u>**Q58699**</u>) which is identified here as containing 14 4Fe-4S clusters (Figure 3b), and a human protein (zinc finger protein 208; UniProt Accession: <u>**O43345**</u>), which is identified here as containing 36 Zn-binding sites (Figure 3c). As large as these numbers might seem, they appear also to be reasonable results: the former protein is annotated in UniProt as containing 13 (not 14) 4Fe-4S-clusters, while the latter is annotated as containing 39 zinc finger domains. Interestingly, three of the latter are classified on the UniProt page as “degenerate” which indicates that they deviate from the established sequence pattern expected of the PROSITE <u>**PUR00042**</u> annotation rule; these are the binding sites that are not identified the AlphaFold2 structure.

**Figure 3.**
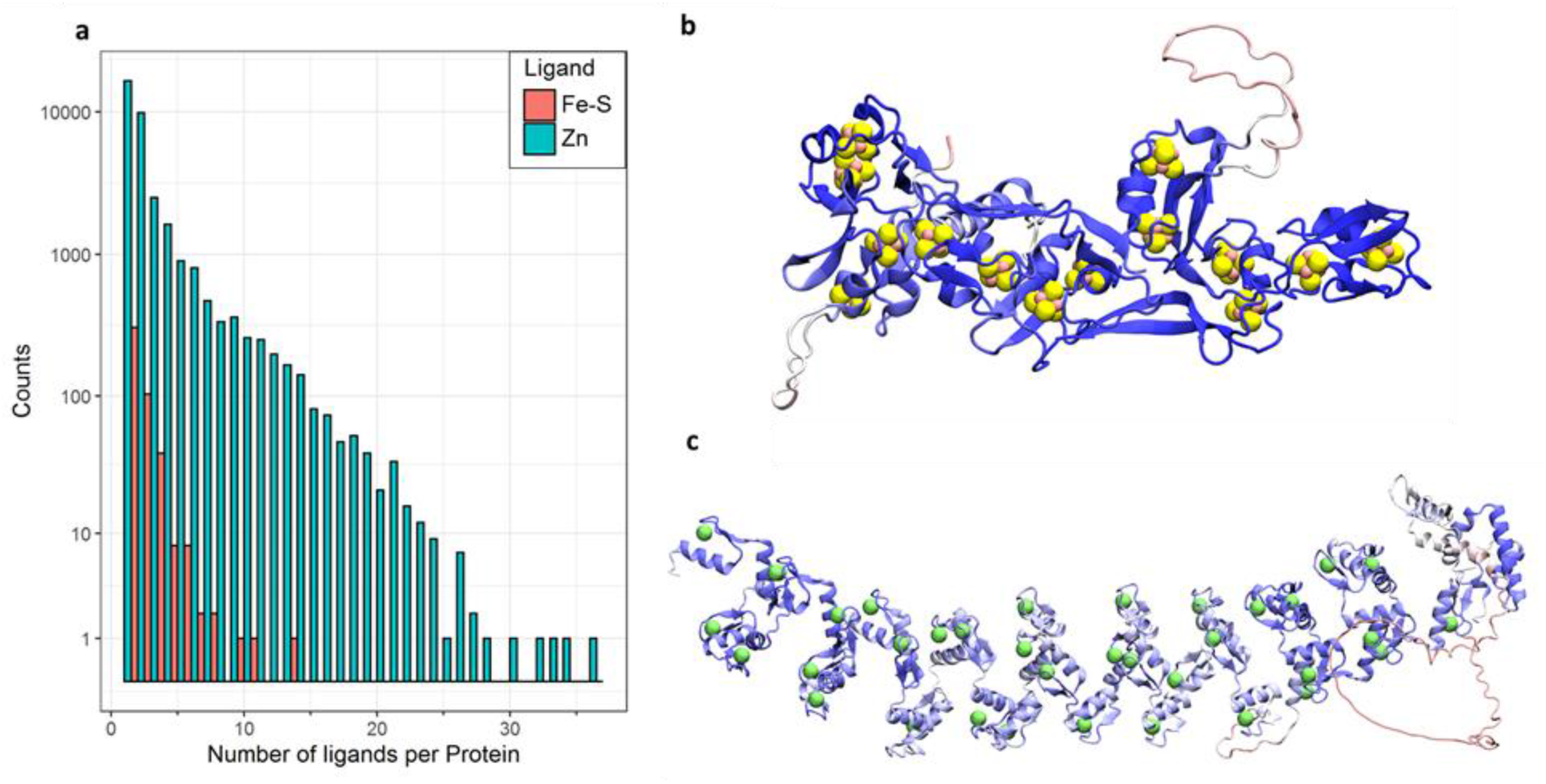
(a) histograms of the number of identified Fe-S clusters and Zn ions per protein. (b) structural view of UniProt Accession: <u>**Q58699**</u>, which has the largest number of identified Fe-S cluster binding sites. (c) structural view of UniProt Accession: <u>**O43345**</u>, which has the largest number of identified Zn binding sites.

### Tens of thousands of Fe-S cluster- and Zn-binding sites are identified in the AlphaFold2 proteomes

Figures 4a-c show the numbers of Fe-S cluster-binding sites and Zn-binding sites identified in all proteins in all 21 organisms for which the AlphaFold2 team has reported structures; note that in this and subsequent figures, organisms are presented in phylogenetic order (see Methods). Figure 4a shows the total number of binding sites identified, regardless of the implied quality of the AlphaFold2 structure. Figure 4b shows the number of binding sites identified when we restrict attention only to those cases for which all coordinating sidechains have a pLDDT score of greater than 70; according to the DeepMind team, residues with a pLDDT score above this threshold are considered “confident” predictions by AlphaFold2 [8]. Finally, Figure 4c shows the number of binding sites identified when we further restrict attention only to those binding sites for which all coordinating side chains have a pLDDT score of greater than 90; such residues are considered “highly confident” predictions by AlphaFold2 [8]. The total number of Fe-S cluster-binding sites and Zn-binding sites identified across all 21 organisms is 91086, 85684, and 37645 for threshold pLDDT scores of 0, 70, and 90, respectively. Imposing the requirement that all coordinating residues be “confident” predictions, therefore, results in only a small decrease in the total number of identified ligands (6% are filtered out) while imposing the more stringent requirement that all coordinating residues be “highly confident” results in a greater decrease (59% are filtered out). In the remainder of this manuscript we focus on binding sites identified when the pLDDT threshold of 70 is applied, but in the Supporting Information all results are provided.

**Figure 4.**
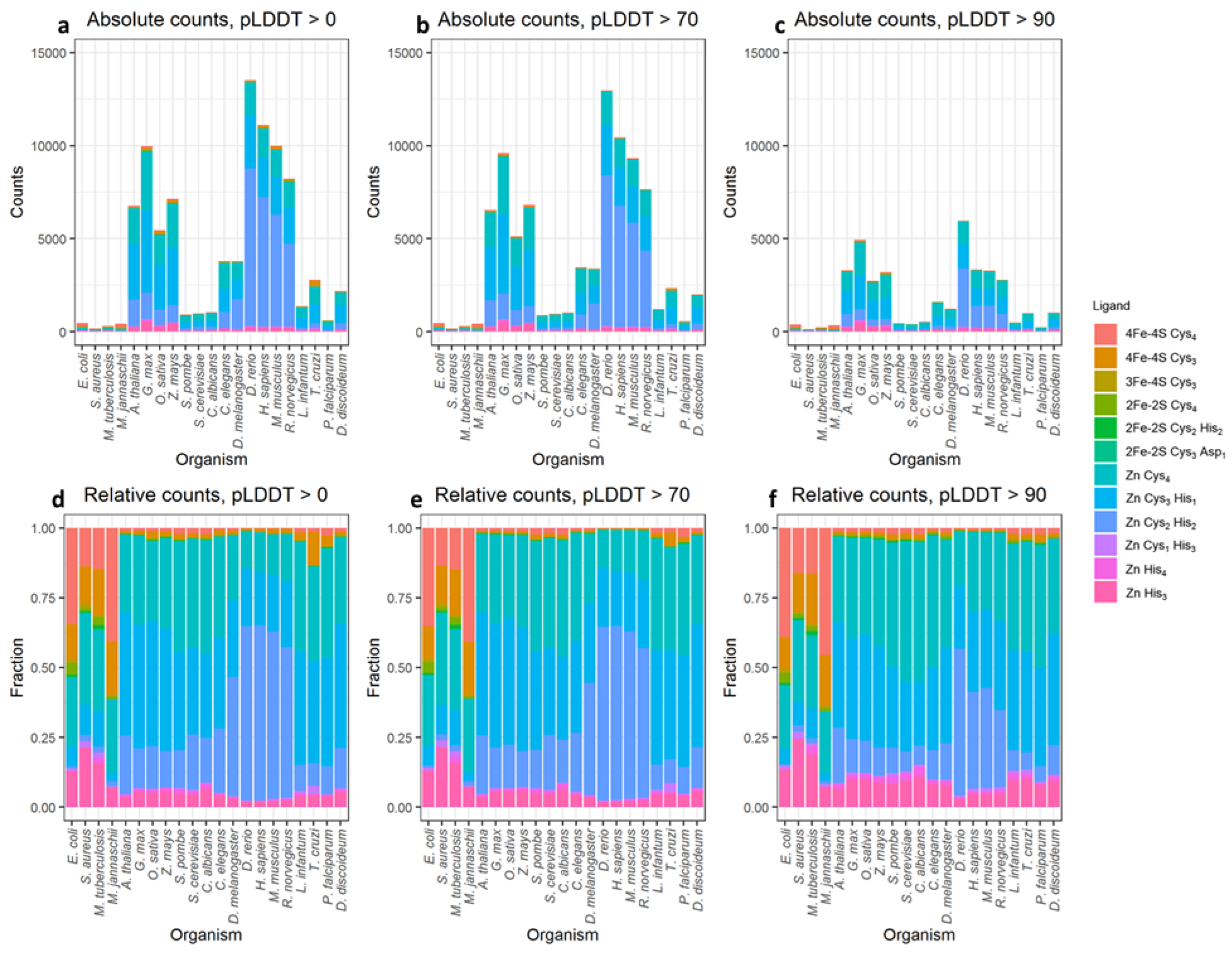
The quantification of identified ligand placements for each ligand type for each organism. (a-c) show the absolute quantification of the identified ligand binding sites, and (d-f) show the relative quantification of the identified ligand binding sites. (a) and (d) show all binding sites, (b) and (e) show binding sites for which all coordinating residues have pLDDT scores greater than 70. (c) and (f) show binding sites for which all coordinating residues have pLDDT scores greater than 90.

The coloring scheme used in Figures 4a-c identifies the total numbers of identified binding sites for all twelve ligand types for which we searched (see Methods). From this it is apparent that we identify far more binding sites for Zn ions than binding sites for Fe-S clusters; this is especially true for the eukaryotic organisms (on the right-hand side of the graphs). In Supporting Information, we show that the higher binding site counts for eukaryotic organisms are not simply due to their proteomes containing more proteins: when we normalize the total count of binding sites by the number of proteins in the proteome of each organism, the eukaryotic organisms again return higher values (Figure S2a). Interestingly, however, if we instead consider the number of proteins predicted to contain one or more Fe-S- or Zn-binding sites and we again normalize by the number of proteins in the proteome, we observe that the difference between the prokaryotes and eukaryotes is much smaller (Figure S2b). In particular, for the 21 organisms, the fraction of proteins within each proteome that we identify as containing at least one Fe-S- or Zn-binding site ranges from 0.05 for *S. aureus* to 0.18 for *D. rerio* (zebrafish). The fraction of Fe-S binding proteins that we identify in *E. coli* is 0.033, which is similar but somewhat lower than a prediction made many years ago that approximately 5% of *E. coli* proteins are Fe-S binding proteins [28].

Finally, we can also examine the relative populations of binding sites identified for each of the twelve ligand types within each organism; these data are plotted using pLDDT thresholds of 0, 70, and 90 in Figures 4d, 4e, and 4f, respectively. From these it can be seen that the relative populations of the various ligand types within each organism show little difference as the pLDDT threshold is increased; this suggests that there is no substantial bias toward building certain ligand binding sites when AlphaFold2’s confidence is low. It can also be seen that the prokaryotic organisms (four left-most columns) are identified here as containing higher relative populations of 4Fe-4S clusters than the eukaryotic organisms, while the eukaryotic organisms contain higher relative populations of Zn-binding sites. Interestingly, a substantial fraction of the 4Fe-4S cluster binding sites identified in the prokaryotes contain 3, not 4 cysteine sidechains (see “4Fe-4S Cys_3_” in Figures 4d-f). Visual examination of these cases indicates that a few of these are “near misses” that contain a fourth potentially coordinating cysteine sidechain nearby (Figure S3a) and that can usually be successfully identified if we loosen the RMSD threshold from 0.50 to 0.75 Å. In most cases, however, there is no fourth obvious coordinating residue but there is instead a vacancy that could be occupied either by a water molecule – as happens, for example, in aconitase (e.g. reviewed in [29]) – or by residues on a separate polypeptide chain not included in the AlphaFold2 prediction (see Discussion); an example of such a case is shown in Figure S3b.

The prokaryotic organisms also appear to have a higher proportion of Zn-binding sites that contain only three coordinating histidine sidechains (“Zn His_3_”). Similar to what was described above, visual examination of these cases reveals that a substantial fraction (26 out of 176 cases) do in fact contain a fourth histidine sidechain but that it points in a non-coordinating direction (e.g. Figure S4a). In a substantial fraction of other cases (36) a clear fourth coordinating sidechain is also present, but it is either an aspartate or glutamate for which we did not explicitly search (e.g. Figure S4b); the prevalence of binding sites with one acidic sidechain is consistent with Zn-binding sites documented in databases [17, 18]. In the majority of cases (114), however, the fourth coordination site is again vacant and solvent exposed (e.g. Figure S4c).

### The relative frequencies of binding sites identified within organisms match the relative frequencies of their annotations in UniProt

An indirect source of support for the present set of results is to compare the relative populations of ligand binding sites identified here with the relative populations already annotated within UniProt. To do this, we make use of UniProt’s “Sites”, “Domain”, “Region” and “Zinc finger” sections to compile separate lists of annotated binding sites for each type of bound ligand, i.e. 4Fe-4S, 3Fe-4S, 2Fe-2S or Zn (see Methods). A comparison of the relative fraction of the binding sites identified within the AlphaFold2 proteomes for these four ligand types (Figure 5a) with those annotated within UniProt (Figure 5b) shows that they are in good agreement, with the higher prevalence of 4Fe-4S clusters in the prokaryotic organisms especially notable in both panels. This good overall agreement suggests that neither the AlphaFold2 structures themselves nor our search algorithm are unduly biased toward building or identifying particular types of binding sites.

**Figure 5.**
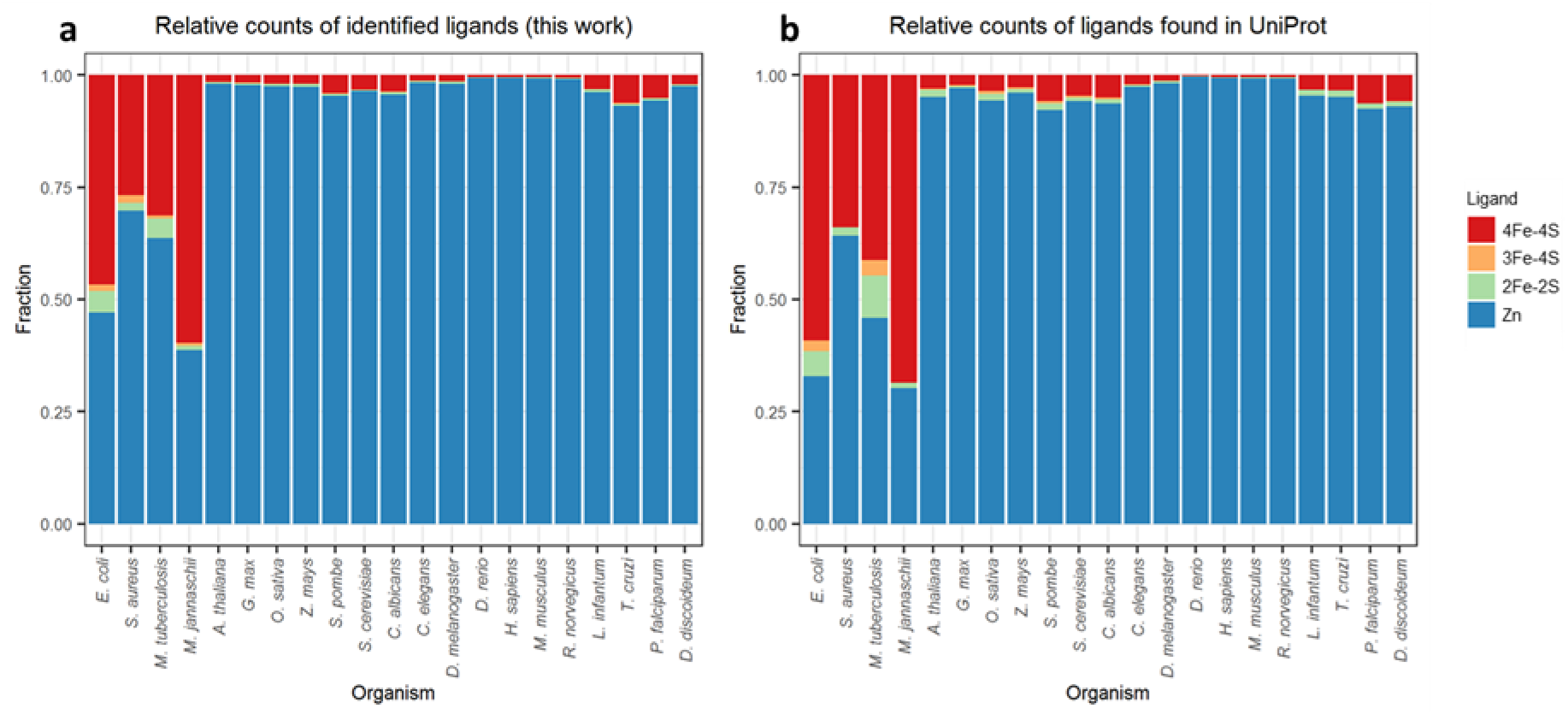
The relative quantification of identified ligand placements for each organism compared to data for the same proteomes extracted from UniProt. (a) relative ligand populations identified here, (b) relative ligand populations extracted from UniProt.

### Recall rates for ligand binding sites annotated within UniProt are high

While the above results suggest that there is little bias in the AlphaFold2 predictions, they do not indicate the extent to which ligand binding sites that are already annotated within UniProt are recovered by our search method. To do this we first compiled a list of all binding sites annotated within UniProt for each of the twelve ligand types for which we searched (see Methods). Those numbers are presented in column three of Table 1, first for the full set of all 21 organisms and then separately for three model organisms (*E. coli*, *S. cerevisiae* and human). We then asked how many of those binding sites were correctly identified by our method – with a correct identification being classed as one in which the ligand and *all* of its coordinating residues were correctly identified; those numbers appear in column four. Finally, we calculated the recall rate as the ratio of these two numbers expressed in percentage form (see column five).

**Table 1.**
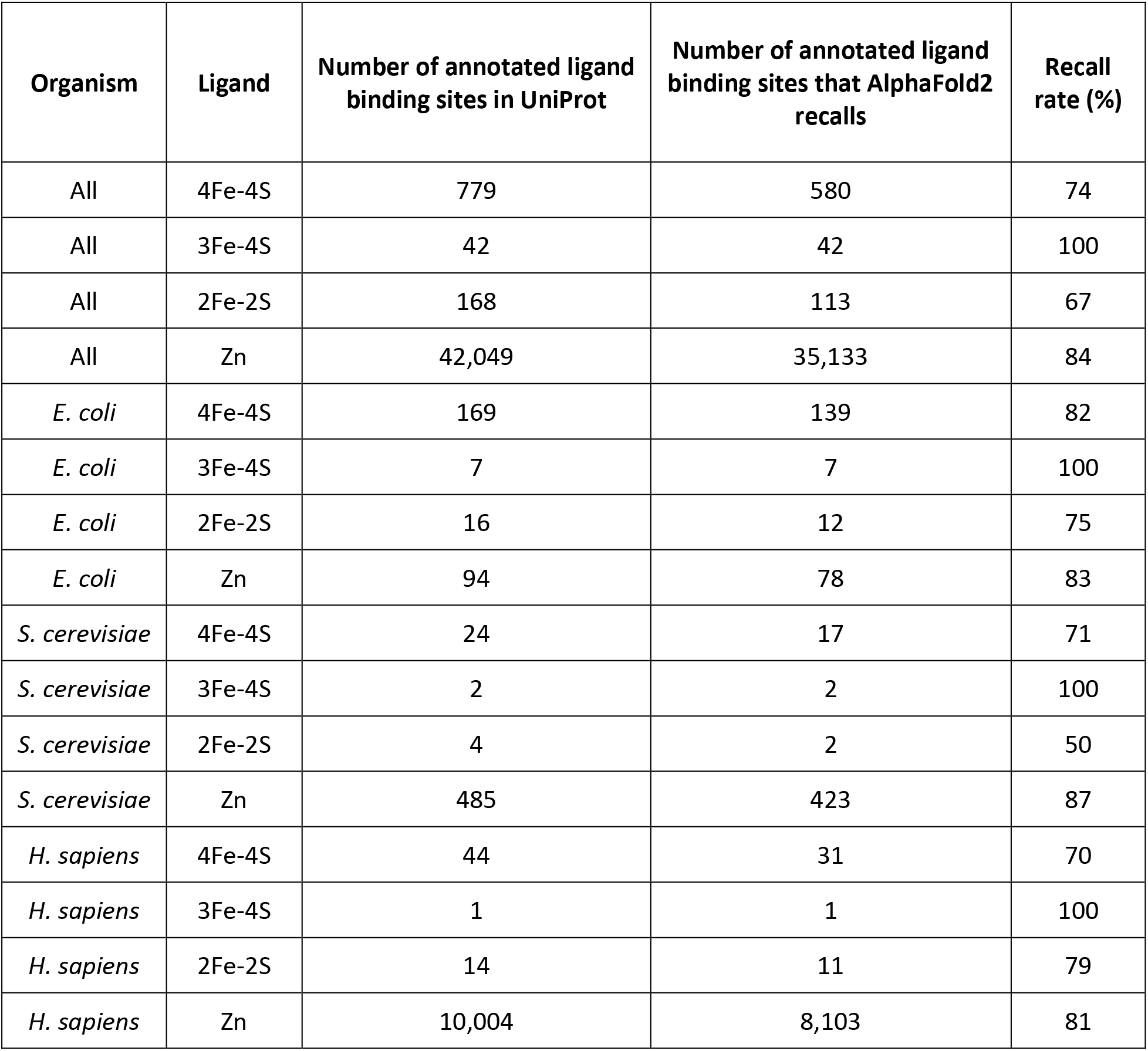

Overall, the recall rates are very good. In the full set of 362,311 proteins, there are 779 4Fe-4S clusters annotated in UniProt that match our search criteria – i.e. “4Fe-4S Cys_4_” or “4Fe-4S Cys_3_” – and 580 of these (i.e. 74%) are correctly identified in the Alphafold2 structures. Visual examination of the 199 4Fe-4S binding sites that are missed by our method suggests that these cases can be classified into five categories. Three of these categories are comparatively uninteresting but are described here to provide a complete accounting: (1) 110 cases are “near misses”; in 102 of these, AlphaFold2 builds a potential binding site with high confidence but the best-fit RMSD for any of our Fe-S ligand types is too high to pass the threshold; in the remaining 8 cases, we can successfully fit “4Fe-4S Cys_3_” but not “4Fe-4S Cys_4_”; (2) 13 cases in which AlphaFold2 built a probably erroneous disulfide bond between coordinating cysteine residues; (3) 11 cases in which AlphaFold2 was unconfident (pLDDT scores less than 70) and did not build the coordinating residues from UniProt into a binding site.

Much more interesting are the remaining two categories, in which AlphaFold2 disputes the binding site information annotated in UniProt. In 45 cases, we obtain what we term “alternative hits”: these are cases in which we identify 4Fe-4S binding sites that share some, but not all, of the UniProt-annotated coordinating residues. In 34 of these “alternative hits”, AlphaFold2 builds 4Fe-4S binding sites by swapping one of the coordinating residues between two different annotated UniProt binding sites within the same protein (e.g. Figure 6a). In the remaining 11 “alternative hits”, AlphaFold2 builds a 4Fe-4S binding site using only three of the four UniProt-annotated coordinating residues and confidently places the fourth coordinating residue well outside of the binding site (e.g. Figure 6b). In the remaining category are 20 cases in which AlphaFold2 rejects completely the UniProt-annotated 4Fe-4S binding site. Importantly, these are not cases in which AlphaFold2 lacks confidence (see above), they are instead cases where AlphaFold2 appears to be confident that there is no 4Fe-4S binding site, and in which it instead often scatters the annotated coordinating cysteines throughout an α helical region of structure (Figure 6c).

**Figure 6.**
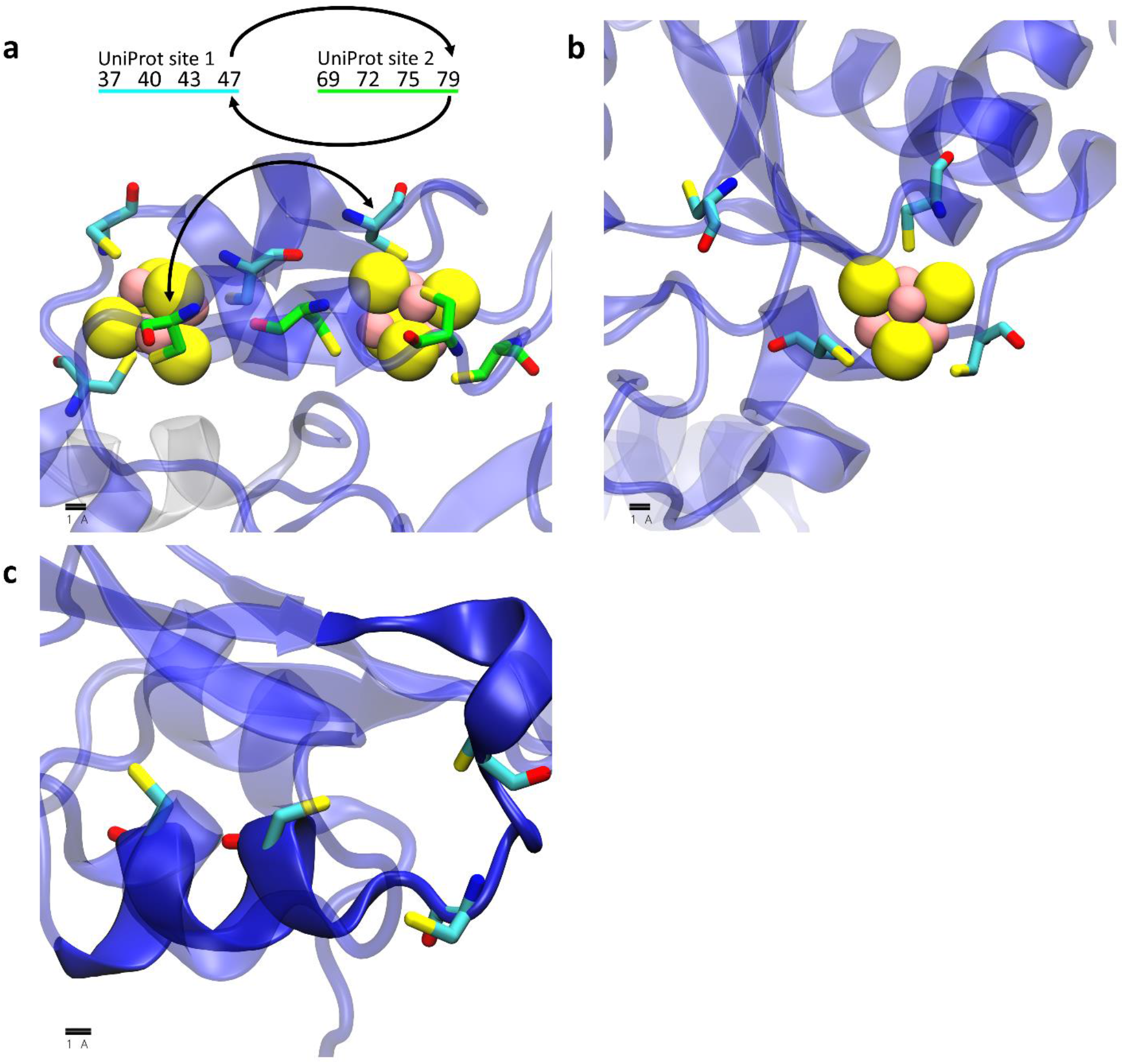
Example 4Fe-4S cluster binding sites annotated in UniProt that are not identified in this work. In each image protein chains are shown as transparent cartoons colored using AlphaFold2’s pLDDT (i.e. confidence) scores on a spectrum of red (low confidence) to blue (high confidence), coordinating residues are shown as licorice, and iron and sulfur atoms in the 4Fe-4S cluster are shown as yellow and pink spheres, respectively. (a) an example in which AlphaFold2 builds two 4Fe-4S sites out of residues annotated in two different UniProt sites but swaps two cysteine residues between each of the sites: the two UniProt sites are shown with their carbon atoms colored in either cyan or green and the residue numbers for the corresponding sites are displayed above the image with arrows to indicate the cysteines that are swapped. (b) an example in which AlphaFold2 confidently builds a “4Fe-4S Cys_4_” site as “4Fe-4S Cys_3_” site and rejects the annotation of the fourth residue. (c) an example in which AlphaFold2 does not build the UniProt-annotated residues into a 4Fe-4S binding site but instead confidently builds the residues in an α helix (opaque blue cartoon).

Good results are obtained for the recall of 3Fe-4S clusters (which have an overlapping definition with 4Fe-4S clusters; see Methods) and for Zn binding sites; 84% of the 43,199 annotated binding sites are correctly identified. Good recall rates for these three types of ligands are also obtained when we consider each of the three model organisms in turn (Table 1). Less good are the results for the 2Fe-2S clusters, for which we recall 113 (i.e. 67%) of the 168 annotated cases. Visual examination of the 55 AlphaFold2 structures that are annotated as containing a 2Fe-2S cluster in UniProt but that are not identified by our method demonstrates that in the majority of cases (34 out of 55), one or more pairs of the cysteines are erroneously disulfide-bonded with each other (Figure 7a). For the other 21 cases, AlphaFold2 builds sidechains that are not quite able to coordinate a 2Fe-2S cluster, and these cases can further be categorized as either those that closely represent a binding site and that are therefore near-misses (Figure 7b) or those with a fourth coordinating residue separated by a great distance from the other three coordinating residues (Figure 7c). Interestingly, one “alternative hit” is identified in the single case of the ferredoxin-like protein from *E. coli* (UniProt: <u>**P0ABW3**</u>): for this protein, Alphafold2 confidently includes a different coordinating residue in the metal binding site (Figure 7d).

**Figure 7.**
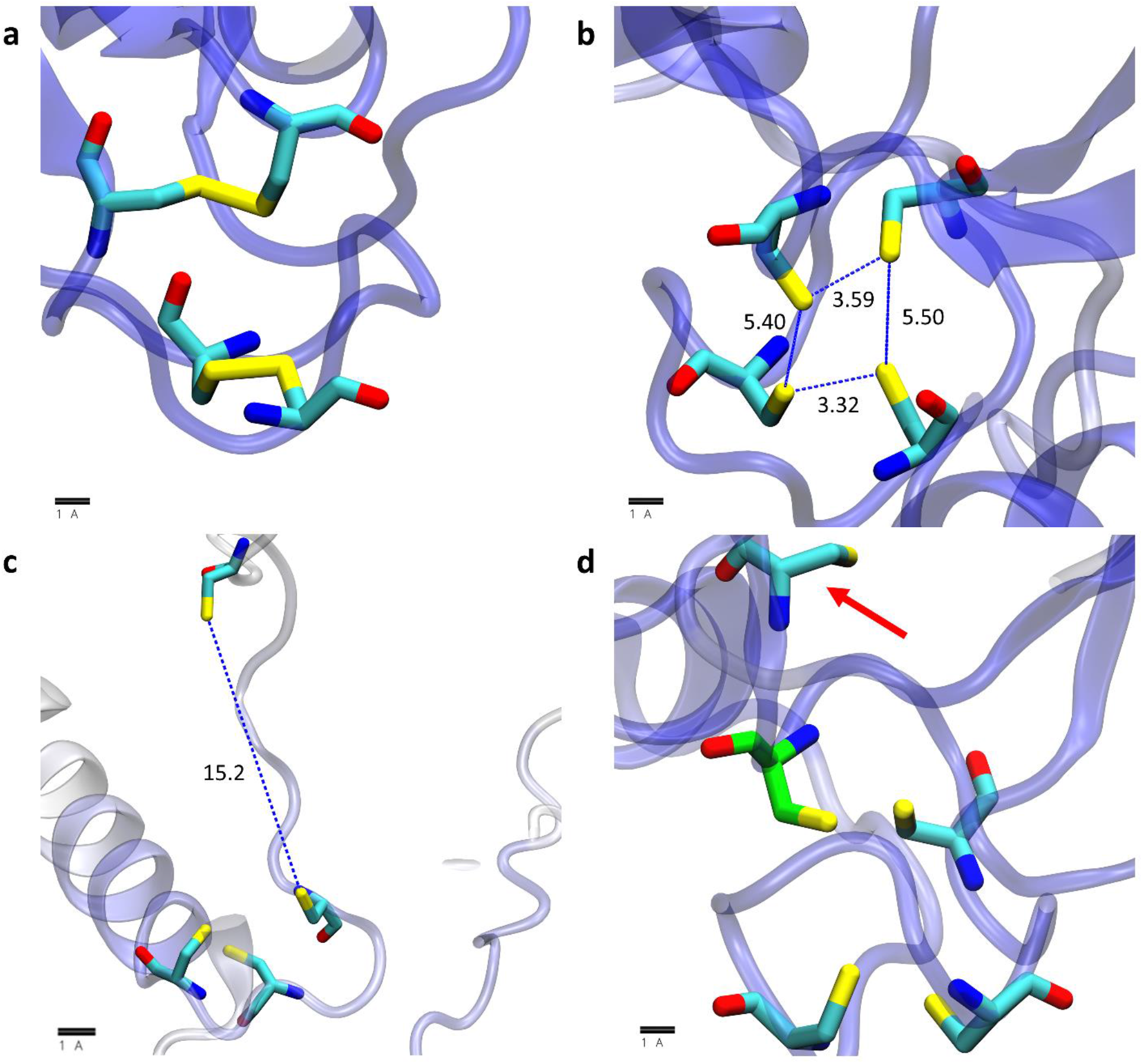
Example 2Fe-2S cluster binding sites annotated in UniProt that are not identified in this work; visualization scheme is the same as used in Figure 6. (a) an example in which AlphaFold2 confidently builds disulfide bonds in the annotated UniProt site. (b) an example in which AlphaFold2 confidently builds a potential site that is too constricted to coordinate a 2Fe-2S cluster. (c) an example in which AlphaFold2 builds the fourth coordinating residue in a position that is more than 5 Å from the three other coordinating residues. (d) an example in which AlphaFold2 builds a potential 2Fe-2S site with a different fourth coordinating residue (carbon atom colored green) than that which is annotated in UniProt (red arrow).

While the above analysis suggests that the lower recall rates for 2Fe-2S clusters relative to those for 4Fe-4S clusters might be mostly attributable to AlphaFold2 occasionally building erroneous disulfide bonds, another possibility that we have considered is whether AlphaFold2’s neural network might be more strongly trained on 4Fe-4S clusters. To explore this issue, we searched through the RCSB to see if 4Fe-4S clusters were represented at much higher levels than 2Fe-2S clusters. At the time of writing, we found 832 structures containing 2Fe-2S (i.e. ligand ID “FES”), 1,392 structures containing 4Fe-4S (i.e. ligand ID “SF4”), and 289 structures containing 3Fe-4S (i.e. ligand ID “F3S”). While these counts include redundant entries (i.e. multiple cases of the same protein), they do not suggest that 2Fe-2S clusters are more poorly represented in the RCSB than 4Fe-4S clusters.

### AlphaFold2-predicted Fe-S cluster binding sites for *E. coli* agree well with previous bioinformatics predictions

For Fe-S clusters, we have also compared the proteins identified here with those identified in two prior studies that developed methods specifically to predict Fe-S cluster-binding proteins. In work reported in 2014, Estellon *et al.* [21] reported successfully identifying 90 out of the 136 Fe-S cluster binding proteins known at the time in *E. coli* using their so-called “mixed” model, and 109 out of 136 using an “extended” model; these recall rates amount to 66 and 80% respectively. In work reported in 2016, Valasatava *et al.* [22] reported successfully identifying 132 out of 149 known Fe-S cluster binding proteins, with a recall rate of 89%. Of the 149 proteins listed in the latter paper 111 are identified here as binding Fe-S-clusters, which represents a recall rate of 74%, i.e. comparable to these previous methods, but somewhat lower. Interestingly, there is extensive overlap between the predictions of all three methods (Figure S5): over 70% of all Fe-S proteins are predicted by each method (*i.e.*, 96 out of 134 Fe-S cluster proteins in our work, 155 from Valasatava *et al.* and 124 from Estellon *et al.*). The binding sites identified here have a noticeably greater degree of overlap with those identified by Valasatava *et al.* (96 + 16 = 112 proteins) than those identified by Estellon *et al.* (96 + 1 = 97 proteins).

We also compared the Fe-S cluster-binding proteins identified here with those that were previously labelled as “false positive” predictions at the time that the works by Estellon *et al.* [21] and Valasatava *et al.* [22] were carried out. In their work, Estellon *et al.* speculated that at least some of their 15 apparent false-positives might be real Fe-S-binding proteins, and they provided experimental evidence implicating two of them as being 4Fe-4S cluster-containing proteins: YdiJ (UniProt ID: <u>**P77748**</u>) and YhcC (UniProt ID: <u>**P0ADW6**</u>). Interestingly, both of those proteins are also successfully identified here, although YdiJ is predicted here to contain two 4Fe-4S clusters and one 2Fe-2S cluster. Of the remaining 13 false-positives predicted by Estellon *et al.*, our method also predicts that four of these proteins contain Fe-S clusters. These predictions are a 4Fe-4S site in YjiM (UniProt ID: <u>**P39384**</u>), a 4Fe-4S site in YhaM (UniProt ID: <u>**P42626**</u>), both a 4Fe-4S and 3Fe-4S site in PreT (UniProt ID: <u>**P76440**</u>) and a 2Fe-2S site in YcbX (UniProt ID: <u>**P75863**</u>), all of which would appear to be excellent candidates for future experimental validation. In their work, Valastava *et al.* produced 23 predictions that were characterized at the time as “false positives”; they noted that homology models of seven of these proteins could be built that contained a plausible 4Fe-4S cluster binding site. Our method predicts 4Fe-4S binding sites in all seven of these proteins in what appear to be similar regions to those shown in images of the homology models reported by Valastava *et al*. For none of the remaining 16 proteins did we find any Fe-S cluster binding sites. In summary, all of these results suggest that, at least in terms of Fe-S cluster-containing proteins, the approach tested here, which is essentially untrained, is comparable in performance to bioinformatics-based prediction methods that have been specifically trained to identify Fe-S cluster binding proteins.

### Tens of thousands of Fe-S cluster binding sites and Zn-binding sites are predicted that are not annotated in UniProt

While it is important to show that AlphaFold2-based predictions can recall annotated cases in UniProt (see above), it is also important to determine the extent to which the present method identifies novel binding sites. To explore this issue, we started with the 85,684 ligand binding sites identified here with a minimum pLDDT score of 70 and eliminated all sites for which there was already a matching record in the corresponding UniProt page (see Methods). After this filtering step, 46,422 of the ligand binding sites (54%) remained; these are the binding sites that we propose might be novel. This number is comparable with the total number of such binding sites already annotated within UniProt for the proteins studied here: 43,031 (some of which we fail to identify); the new binding sites predicted here, therefore, have the potential to roughly double the number of known Fe-S cluster- and Zn-binding sites. In terms of protein counts, we predict an additional 22,560 proteins as being binders of Fe-S clusters and/or Zn ions that were not previously annotated as binders of these ligands in UniProt. Since the number of proteins already annotated in UniProt is 13,094 the new binding sites predicted here have the potential to roughly triple the number of known Fe-S cluster- and Zn-binding proteins. The relative gains for each organism are plotted for novel sites and for novel Fe-S cluster- and/or Zn-binding proteins in Figures 8a and 8b, respectively. Interestingly, both plots show a greater than 6-fold increase for several less extensively annotated organisms (e.g. *G. max*, *Z. mays*, and *T. cruzi*, etc.), but even for the much better characterized human proteome a substantial number of new proteins are found: prior to the current work, 2144 human proteins were annotated in UniProt as Fe-S cluster- and/or Zn-binding proteins; here we find an additional 638 proteins, thus representing an increase of 29%.

**Figure 8.**
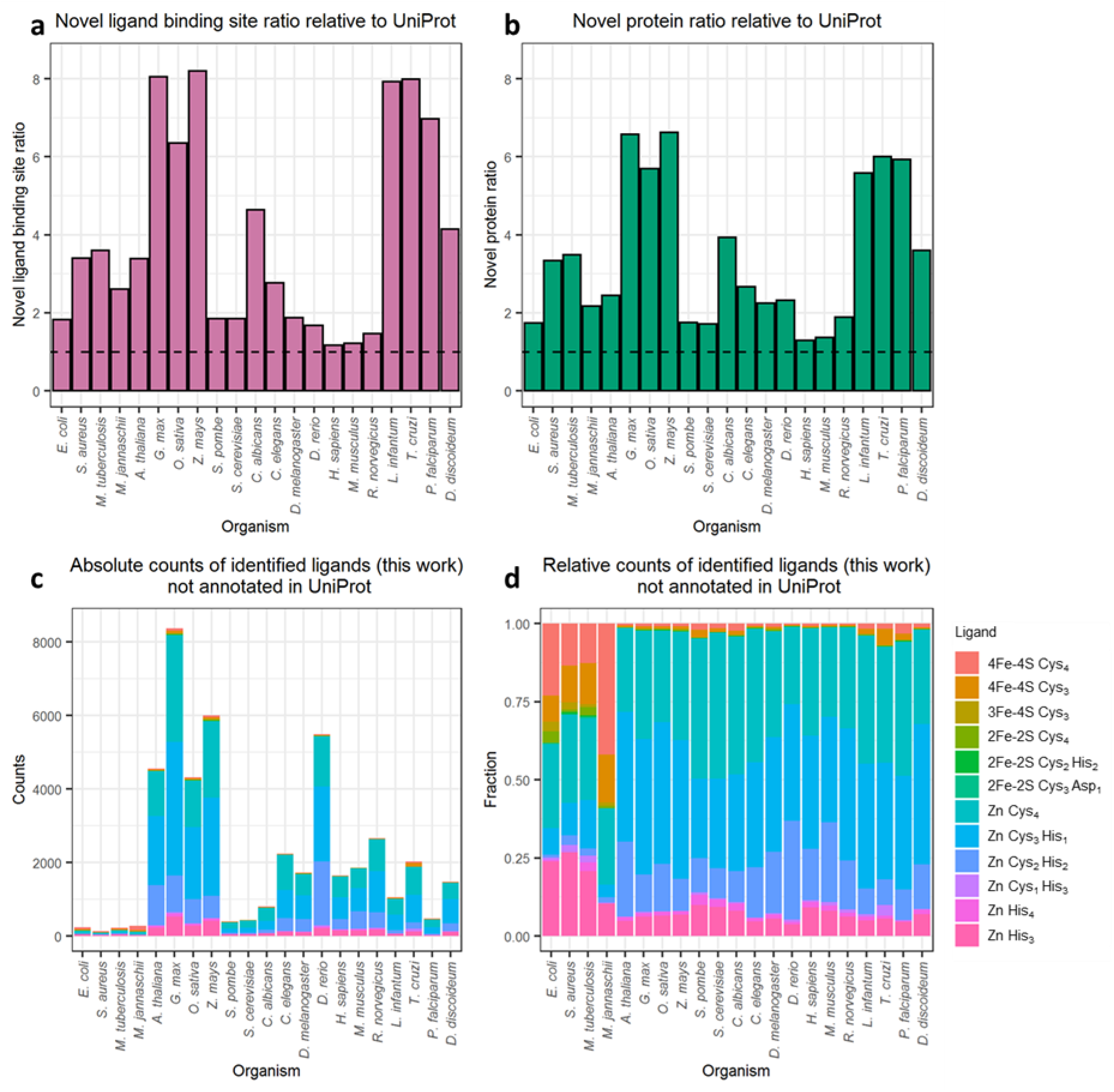
Quantification of novel ligand binding site predictions with pLDDT scores greater than 70 and not annotated in UniProt. (a) the ratio of ligand binding sites identified here to those already annotated in UniProt for each organism. (b) the ratio of ligand binding proteins identified here to those already annotated in UniProt for each organism. For both panels the dashed line denotes a ratio of 1, which indicates no increase in the size of the metalloproteome of a given organism. (c) absolute counts of proposed novel binding sites for each ligand type in each organism. (d) relative counts of proposed novel binding sites for each ligand type in each organism.

Figure 8c shows that the vast majority of the proposed new binding sites in eukaryotic organisms are Zn binding sites. Figure 8d, which is a normalized version of Figure 8c, demonstrates the increased relative abundance of Fe-S clusters in the novel predictions in prokaryotic organisms (four left-most columns). Importantly, the distributions of RMSDs for identified ligand binding sites already present in UniProt (Figure S6a) and for identified ligand binding sites not already present in UniProt (Figure S6b) are very similar. This indicates that the quality of the predictions and their resulting reliability is indistinguishable from the known cases.

### Thousands of Fe-S and Zn-binding sites are predicted in proteins with no structural homolog

While the above analysis identifies and highlights those binding sites that are not already explicitly annotated in UniProt, it could be argued that many such sites might be rather easy predictions for AlphaFold2 to make if the protein has a homologous structure already in the RCSB. In order to be as strict as possible in labeling a binding site as a truly novel prediction, therefore, we applied an additional filter that eliminated those binding sites for which a remote homolog was identifiable in the PDB70 database [30] that aligned with *any* of the predicted coordinating residues (see Methods). With this extra, and very stringent filtering step, 13,139 binding sites predictions in 7,490 unique proteins remain as novel predictions. The relative gains for each organism are plotted for novel ligand binding sites and proteins in Figures 9a and 9b, respectively; as expected, the addition of the hhsearch filter, results in smaller relative gains (compare Figures 8a/b with Figures 9a/b, respectively). Figure 9 c-d recapitulates most of the trends seen in Figures 8 c-d with the predicted Zn binding sites again dominating both the absolute and relative counts. As above, the distributions of RMSDs for predictions that have a remote homolog or are found in UniProt (Figure S7a) are very similar to those obtained for the predictions that have no remote homolog and that are not found in UniProt (Figure S7b). This suggests that the ligand binding sites built by AlphaFold2 for which there is no close structural homolog in PDB70 have similar quality to those that do.

**Figure 9.**
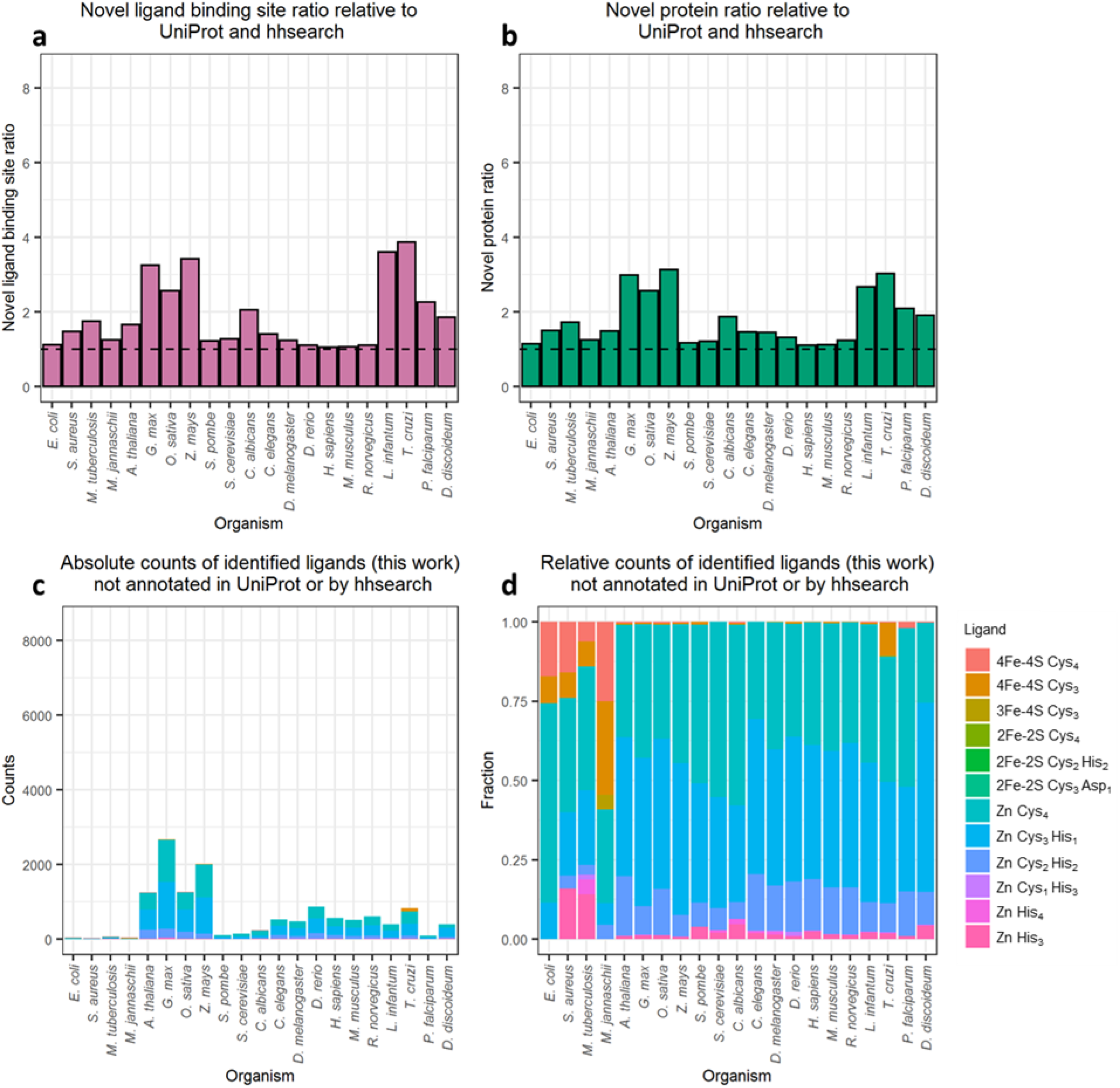
Quantification of novel ligand binding site predictions with pLDDT scores greater than 70, not annotated in UniProt, and without a remote homolog identified by hhsearch. (a) the ratio of ligand binding sites identified here to those already annotated in UniProt for each organism. (b) the ratio of ligand binding proteins identified here to those already annotated in UniProt for each organism. (c) absolute counts of proposed novel binding sites for each ligand type in each organism. (d) relative counts of proposed novel binding sites for each ligand type in each organism.

### Cysteines predicted here to be part of Fe-S cluster or Zn-binding sites generally have lower chemical reactivities *in vivo*

Many of the Fe-S cluster-binding sites and Zn-binding sites predicted here must await experimental verification. In lieu of that, one test that we can immediately perform is to determine the extent to which cysteine residues predicted here to be parts of ligand binding sites are found to be chemically unreactive in chemical proteomics experiments carried out on living cells (e.g. [31]). Recent isoTOP-ABPP experiments, for example, have probed the chemical reactivity of cysteine residues on proteomic scales in human cancer cell lines [32] and in *E. coli* [23]. Given that cysteine residues identified experimentally as “highly reactive” are generally assumed to be unlikely to permanently coordinate ligands such as Fe-S clusters or metal ions (e.g. [33]), we should expect to find little overlap between cysteines that we identify here as being part of ligand binding sites and cysteines found to be highly reactive in isoTOP-ABPP experiments.

To explore this point, we performed separate analyses on human and *E. coli* proteins, comparing with the combined dataset of three different cancer cell lines for the former [32], and using the single dataset for the latter [23]. The results for the human proteins are shown in Figure 10a. Encouragingly, the left panel shows that the subset of highly reactive cysteines (blue circles) and the subset of ligand-coordinating cysteines identified here (red circles) overlap with each other much less than is expected by chance (234 common residues; p-value = 5.41e-17 according to a one-tailed Fisher’s exact test). In contrast, and as hoped, the middle panel of Figure 10a shows that the subset of ligand-coordinating cysteines identified here (red circles) overlaps with the subset of UniProt-annotated cysteines (grey circles) much more than is expected by chance (538 common residues; p-value < 1e-300). Both of these results are exactly as expected if the predicted ligand binding sites identified in the AlphaFold2 structures are realistic. Finally, the right panel of Figure 10a shows that the subset of highly reactive cysteines (blue circles) overlaps with the subset of UniProt-annotated cysteines (grey circle) slightly less than is expected by chance (65 common residues; p-value = 0.340).

**Figure 10.**
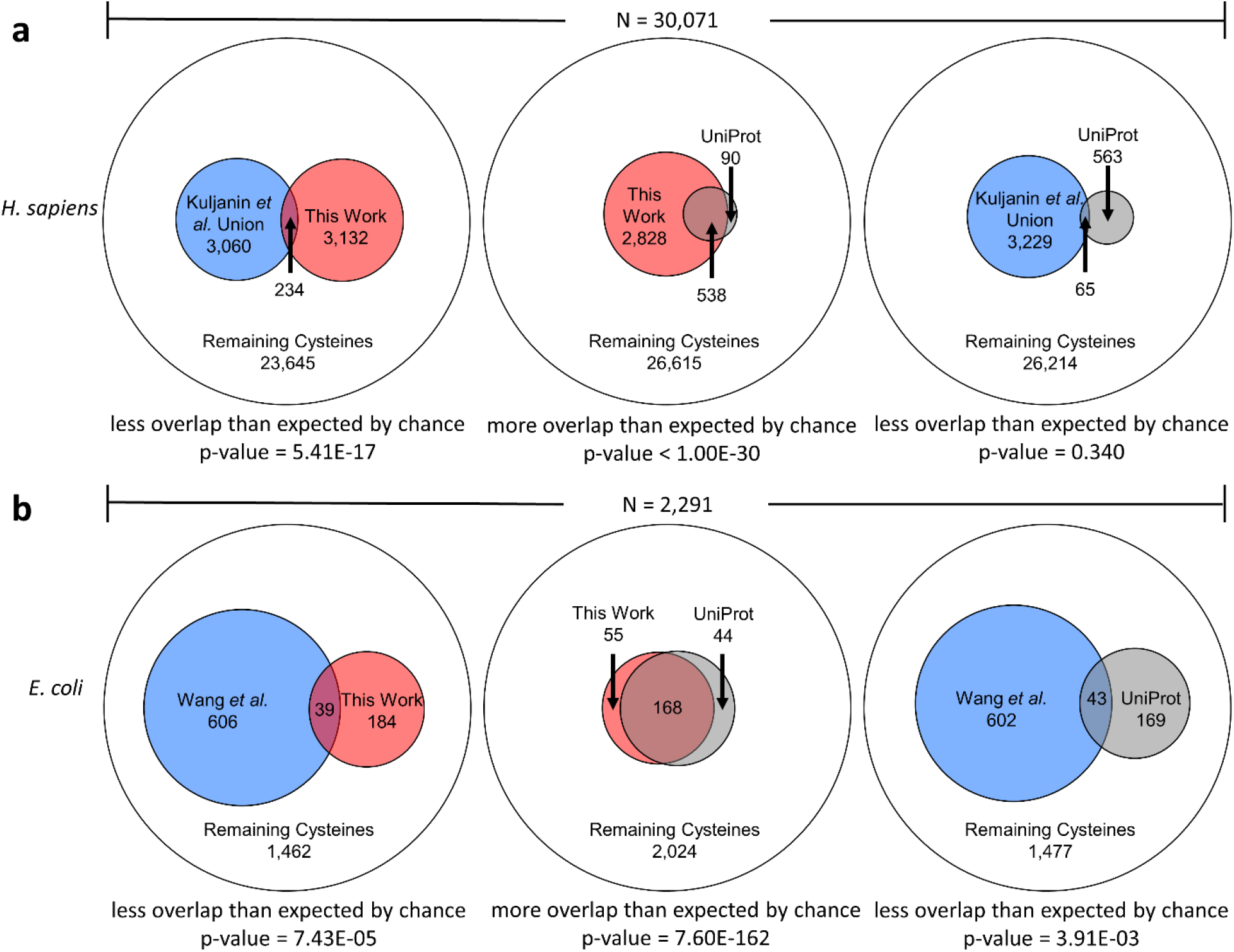
Overlap between subsets of cysteines that are: (1) highly chemically reactive in experiment (blue), (2) identified here as members of ligand-binding sites (red) and (3) annotated in UniProt as members of ligand-binding sites (grey). Comparisons are represented as Euler diagrams: the outer circle represents the total number of cysteines in proteins that contained chemically reactive cysteines in experiment. p-values from one-tailed Fisher’s Exact Tests are shown below each diagram, together with the nature of the hypothesis being tested. (a) shows comparisons for the union of three *H. sapiens* cancer cell lines reported by Kuljanin *et al.* [32], (b) shows comparisons for the *E. coli* data reported by Wang *et al.* [23].

Figure 10b shows corresponding results for *E. coli*. Again, the left panel of Figure 10b shows that the subset of highly reactive cysteines (blue circles) and the subset of ligand-coordinating cysteines identified here (red circles) overlap with each other much less than is expected by chance (39 common residues; p-value = 7.43e-05). Again, in contrast, the middle panel of Figure 10b shows that the subset of ligand-coordinating cysteines identified here (red circles) overlaps with the subset of UniProt-annotated cysteines (grey circles) much more than is expected by chance (168 common residues; p-value = 7.60e-162). Finally, the right panel of Figure 10b shows that the subset of highly reactive cysteines (blue circles) overlaps with the subset of UniProt-annotated cysteines (grey circle) less than is expected by chance (43 common residues; p-value = 3.91e-03).

For completeness, we note that all of the relationships described above are qualitatively independent of our applied pLDDT threshold (i.e., pLDDT > 70) or the definition of “highly reactive cysteines” used by the experimental authors for either *H. sapiens* (Figure S8) or *E. coli* (Figure S9).

### Predicted binding sites are usually conserved between multiple AlphaFold2 fragment models

For 20 of the 21 proteomes studied here, no AlphaFold2 prediction has been reported for proteins exceeding 2700 residues in length owing to their much greater computational requirements. For the human proteome, however, a series of fragment predictions have been reported for 208 proteins ranging in size up to 34,923 residues (titin); each of these fragments contains up to 1400 residues, with each one offset by 200 residues from the preceding fragment. Encouragingly, of the ligand binding sites identified here that reside within these 208 proteins, 178 appear in more than one fragment model, with 33 of the identified binding sites appearing in all 7 fragments that encompass the binding site’s coordinating residues (the maximum possible), and a further 31 of the identified binding sites appearing in 6 fragments. These results suggest that binding sites built by AlphaFold2 in one fragment tend to be conserved within fragments that contain the same sets of coordinating residues. Most interestingly, we found only one protein for which fragment structures contained fundamentally different binding sites, and this was because one of the fragments was missing some of the coordinating residues. Specifically, for the protein (<u>**O43149**</u>) a Zn Cys_2_ His_2_ binding site was confidently built by AlphaFold2 in all six fragments that contain all four coordinating residues (Cys1797, Cys1800, His1819, His1823). In a seventh fragment that contained the two cysteines but not the two histidines, AlphaFold2 instead built a Zn Cys_4_ binding site (adding Cys1783 and Cys1786), but it did so with very low confidence (Figure S10).

## Discussion

In this work we have sought to determine the extent to which Fe-S cluster and Zn binding sites can be identified within the huge dataset of new protein structures made available by the DeepMind team [9]. The results indicate that AlphaFold2 routinely builds high quality binding sites that are specific for one particular type of ligand, and that the placement of the ligand within the binding site is essentially unambiguous and free of egregious steric clashes. The latter two aspects are especially remarkable given that the AlphaFold2 predictions have been made in the absence of any explicit information regarding potential ligands.

The results presented here also demonstrate that AlphaFold2 – even without any explicit training for this purpose – has an ability to recall known Fe-S cluster binding proteins in *E. coli* that appears to be competitive with previous explicitly-trained bioinformatics-based methods. AlphaFold2 would, therefore, seem to be a useful complement to experimental chemoproteomics techniques that attempt to identify metal binding proteins on a proteomic scale. This idea is supported by the fact that, for human and *E. coli* proteins, cysteines identified here to be involved in Fe-S cluster and Zn binding tend to be less chemically reactive than other cysteines in the same proteins (Figure 10). One interesting additional comparison that we can make at the time of writing is with very recent work reported by Bak and Weerapana [15] who have developed a chemoproteomic strategy that can identify 94 out of 144 previously annotated Fe-S-containing proteins in *E. coli* while also predicting a further 14 proteins as potential Fe-S cluster-containing proteins. Two of these 14 proteins (TrhP (<u>**P76403**</u>) and DppF (<u>**P37313**</u>)) were explicitly tested and confirmed by the authors, and encouragingly, both of these proteins are also identified here as binding Fe-S clusters: Trhp is identified here as containing a 4Fe-4S cluster which agrees with the assignment made by Bak and Weerapana, albeit only when we increase the RMSD threshold from 0.50 to 0.56 Å, and DppF is identified as containing a 4Fe-4S cluster, although with different coordinating residues than the single cysteine implicated experimentally [15]. No Fe-S cluster binding sites were identified here in the remaining 12 proteins listed by Bak and Weerapana. While directed experimental methods should remain the gold standard for determining that a protein is capable of binding Fe-S clusters, if confirmed, these latter results would indicate that computational predictions based on AlphaFold2 structures could, at the least, help to prioritize candidate proteins identified in proteome-scale experiments for further study.

The identification here of tens of thousands of potential new Fe-S cluster and Zn binding sites, dramatically increasing the size of the known metalloproteome across twenty-one organisms. Importantly, we have shown that the apparent quality of the proposed novel binding sites is essentially indistinguishable from the quality of the known binding sites, regardless of whether one requires the definition of “novel” to include or exclude sites that have remote structural homologs in the PDB (see Figures S6 and S7). As might be anticipated, AlphaFold2 predicts proportionally greater numbers of new binding sites for organisms that are less well-annotated in UniProt. This suggests a possible role for AlphaFold2 in aiding the functional annotation of unstudied proteomes, especially since in a number of cases it makes predictions that diverge significantly from those already annotated in UniProt. While there are certainly instances where AlphaFold2 appears to make mistakes (see below), in other cases the “alternative hits” predicted with different binding site compositions appear to be sufficiently plausible that it seems reasonable to call the current UniProt annotation into question.

There are two general limitations of the predicted structures made available by the DeepMind team that have a small impact on our ability to identify Fe-S cluster and Zn binding sites. The first limitation is that all of the predicted structures represent proteins in their monomeric state. This prevents the method presented here from reliably identifying those Fe-S clusters and Zn-binding sites whose coordinating residues might be shared between different polypeptide chains. There are, for example, known cases of Fe-S clusters whose binding sites are formed by homo-oligomeric interactions (e.g. [34]). As this manuscript is being completed rapid advances are being made in adapting and extending AlphaFold2 to allow it to model homomeric and heteromeric protein complexes (e.g. [35–37]). It seems likely, therefore, that in the near future the same strategy reported here might be used to identify ligand binding sites that lie at the interface between separate polypeptide chains. The second, more minor limitation for the present application is that, with the exception of the human proteome, structures of very large proteins (i.e. those exceeding 2700 residues) have not been reported. For the human proteome, the DeepMind team has implemented a workaround that models larger proteins using a series of overlapping fragments, each of which represents at most 1400 residues. Encouragingly, we have found here that in many cases the same potential binding sites, employing the same coordinating residues, are predicted in multiple fragments of the same protein (see Results). This suggests that, if the computational expense of modeling very large proteins as single entities proves difficult to overcome, then the alternative approach of modeling them as overlapping fragments is likely to be a good one.

Aside from these general limitations, we have uncovered one scenario in which AlphaFold2 appears prone to making occasional mistakes. Binding sites for Fe-S clusters and Zn ions often involve significant numbers of cysteine residues brought into close proximity. In a number of cases that we have visually examined we have noticed a tendency for disulfide bonds to be added between cysteines that ought, instead, to coordinate a Fe-S cluster or a Zn ion. We have seen this occur in Zn Cys_4_ binding sites, and in binding sites for 4Fe-4S and 2Fe-2S clusters; the fact that it occurs more frequently with 2Fe-2S binding sites than with 4Fe-4S binding sites is likely due to the fact that pairs of Cys SG atoms are closer in the former (^~^3.5 Å) than in the latter (^~^6.3 Å). While excessive disulfide bonding does appear to occur, it is probably important not to over-emphasize this result. It is, after all, extraordinary that AlphaFold2 can build highly plausible binding sites even in the absence of the ligand, and cases where binding sites are malformed due to the presence of erroneous disulfide bonds should be relatively easy to identify.

In closing, we note that the proposed atomic coordinates for all of the bound ligands identified here can be found in Supporting Information. Addition of the coordinates of the proposed Fe-S clusters to the original AlphaFold2 structures is likely to be especially important for those interested in performing virtual screening to identify druggable sites within the AlphaFold2 structures; if not added, screening efforts might be misled by the apparent presence of a cavity that, in reality, is likely already occupied. Finally, we note that while the ligands studied here are all small, the success achieved here would suggest that conducting similar binding site searches for larger, more complex ligands might be a worthwhile undertaking.

## Materials and Methods

### Protocol for identifying potential ligand binding sites in AlphaFold2 structures

The work reported here makes use of code written predominantly in-house that attempts to find potential binding sites within an AlphaFold2 protein structure given a user-specified list of ligand types, together with representative structures of each ligand type (see below). Here we use the phrase “ligand type” to mean a specific type of ligand (e.g. a 4Fe-4S cluster, a single Zn ion, etc) together with a specified set of coordinating residues (e.g. 4 cysteines, 3 cysteines + 1 histidine etc).

The procedure (Figure 1) starts with reading the AlphaFold2 pdb file and continues as follows. A list of all sidechain or backbone atoms that could potentially coordinate one of the input ligand types is made. For those ligand types for which a coordinating residue is a histidine sidechain we select both ND1 and NE2 atoms as potential binding sites within the AlphaFold2 structure; for those ligand types for which a coordinating residue is an aspartate sidechain we select both OD1 and OD2 as potential binding sites. The full set of selected potential coordinating atoms is then clustered into “regions” using a standard single-linkage clustering algorithm with a distance threshold of 8 Å. Each region within the protein is then analyzed in turn. Each ligand type is superimposed at all possible locations within each region; this is achieved by cycling through all possible ways by which the ligand type’s coordinating atoms can be paired with corresponding atoms in the region. In some cases, there can be many possible superpositions to consider. For example, when we attempt to place the 4Fe-4S ligand type with 4 coordinating cysteine sidechains (ligand type: “4Fe-4S Cys_4_”) within a region that contains nine cysteine sidechains, there are, in principle, 9! / (5! 4!) = 126 possible combinations of atoms that could be used to perform the superposition of the ligand within the region. All such combinations are attempted, and those for which the root-mean-squared deviation (RMSD) of the superposing atoms is above a specified threshold (0.5 Å for most of the results reported here) are immediately rejected. Combinations for which the RMSD is below the specified threshold are then checked for steric clashes, and combinations in which any atoms of the ligand are within 2 Å of protein atoms are rejected, as are ones in which any atoms of the ligand are within 2.5 Å of any previously placed ligand. The more lenient value of 2 Å is used for evaluating ligand-protein clashes in an attempt to account for the fact that AlphaFold2’s “predictions” of ligand binding sites are made in the absence of the ligand and so do not always place atoms in a perfect arrangement for accommodating ligands. Having rejected all combinations with poor RMSDs or steric clashes, the remaining combinations are ordered by RMSD and the one with the lowest RMSD is retained. This procedure is carried out for all listed ligand types (12 in the present study). If multiple ligand types are found to have at least one combination that satisfies the RMSD and clash criteria defined above then the ligand type having the highest priority (i.e. the one appearing first in the list of ligand types provided by the user) is retained and the ligand is placed into the structure. The entire superposition process is then repeated for all listed ligand types within the same region until no further ligand type can be successfully placed within it. At that point, the code proceeds to analyze the next region.

Note that during the addition of ligands to the structure, each cysteine sidechain is allowed to coordinate up to two different ligand types: this allows the code, for example, to successfully identify cases where two or more Zn ions are simultaneously coordinated by cysteine sidechains to form a cluster that effectively has a stoichiometry such as Zn_2_ Cys_6_ or Zn_3_ Cys_8_ (see, for example, Figure 1c).

### Selection of representative structures for each ligand type

In the present work we have compiled a list of twelve ligand types for which we have searched for binding sites within the AlphaFold2 structures. Prototypical structures of each of these ligand types have been selected directly from the RCSB, with the specific example chosen being the one with the highest resolution structure in the most-populated sub-category for each ligand in the comprehensive MetalPDB database [17]. For each selected structure we save only the coordinates of the ligand atoms and the directly-coordinating atoms of the coordinating residues (e.g., the “SG” atom in cysteine).

Six of the twelve ligand types that we define involve the common Fe-S clusters. For 4Fe-4S clusters, we define two possible ligand types: (1) a 4Fe-4S cluster with four coordinating cysteines (“4Fe-4S Cys_4_”; from pdb code 3A38), and (2) the same cluster but with one of the four cysteines removed (“4Fe-4S Cys_3_”). This second ligand type is added to allow us to identify 4Fe-4S clusters for which the fourth coordinating atom comes from a residue other than cysteine (e.g. glutamate), or from an S-Ado-Met (e.g. [38]), or from a water molecule (e.g. [39]). For 3Fe-4S clusters, we define one possible ligand type with three coordinating cysteines (“3Fe-4S Cys_3_”; taken from 1WUI). Note that with our procedure this ligand type is usually impossible to distinguish from the 4Fe-4S cluster with three coordinating cysteines, so higher priority is given to the 4Fe-4S cluster. For 2Fe-2S clusters, we define three possible ligand types: (1) a 2Fe-2S cluster with four coordinating cysteines (“2Fe-2S Cys_4_”; taken from 1N62), (2) a 2Fe-2S cluster with two coordinating cysteines and two coordinating histidines (“2Fe-2S Cys_2_ His_2_”; from 3D89), and (3) a 2Fe-2S cluster with three coordinating cysteines and one coordinating aspartate (“2Fe-2S Cys_3_ Asp_1_”; from 1NEK). While not exhaustive, these choices appear to cover the most common sets of coordinating residues found in Fe-S clusters (e.g. [40]).

The remaining six ligand types for which we search involve various combinations of a Zn ion surrounded by three to four coordinating residues. Accordingly, we refer to each of the following as potential Zn binding sites. For such cases we define the following ligand types: (1) a Zn ion with four coordinating cysteines (“Zn Cys_4_”; from 2PVE), (2) a Zn ion with three coordinating cysteines and one histidine (“Zn Cys_3_ His_1_”; from 3SU6), (3) a Zn ion with two coordinating cysteines and two histidines (“Zn Cys_2_ His_2_”; from 4EGU), (4) a Zn ion with one coordinating cysteine and three histidines (“Zn Cys_1_ His_3_”; from 5UAM), (5) a Zn ion with four coordinating histidines (“Zn His_4_”; from 3TIO), and (6) a Zn ion with one of the four coordinating histidines removed (“Zn His_3_”; from 5UAM). The latter was added because it is one of the most common Zn binding sites listed in ZincBind, but as was the case with the “4Fe-4S Cys_3_” ligand type, it also allows us to find cases of binding sites that involve a fourth coordinating residue that is not cysteine. Note that our choice of ligand types for Zn ions reflects the fact that the vast majority of Zn-binding sites documented in MetalPDB [17] and ZincBind [18] involve coordination by only cysteine and histidine, so our choices cover every possible combination of Zn-Cys_x_ His_y_ binding sites, where x and y sum to four.

We note that while we search explicitly for binding sites for Zn ions coordinated by four cysteine sidechains (“Zn Cys_4_”), in rare cases it appears possible that such binding sites might instead be for Fe ions. Literature estimates for the average bond length connecting Fe ions to the SG atoms of their coordinating cysteines average 2.26 Å; [41]), while those connecting Zn ions average 2.34 ± 0.12 Å; [42]). These values are sufficiently similar that given the expected resolution of the AlphaFold2 structures it appears to be effectively impossible to distinguish between them. We expect the vast majority of cases to be genuine Zn binding sites given that there are 1610 Zn binding sites explicitly annotated in UniProt for the 21 proteomes under consideration here, and only 8 such Fe binding sites. Nevertheless, it should be remembered that it is faintly possible that some of the binding sites that we identify here as binding sites for “Zn Cys_4_” might instead be binding sites for “Fe Cys_4_”.

As noted above, when the ligand superposition procedure finds that multiple ligand types can successfully fit into the same “region” of the structure, priority is given to the ligand type that appears first in the input list. An extraordinary feature of the AlphaFold2 structures is that, when ligand binding sites are present, they appear to be highly specific for a particular type of ligand (see Results); given that we also generally explicitly exclude cases where the RMSD of the coordinating atoms exceeds 0.5 Å, there is effectively zero probability of finding cases where the same sets of coordinating atoms within a region are predicted to be binding sites for two quite different ligand types. Instead, the cases where multiple ligand types successfully fit involve ligand types that are structurally almost indistinguishable from each other: e.g. “4Fe-4S Cys_4_” versus “4Fe-4S Cys_3_” versus “3Fe-4S Cys_3_”, and “Zn Cys_4_” versus “Zn Cys_3_”. For each of these cases we assign priority in the order listed, i.e. with preference given to the ligand type that has the greater number of coordinating residues.

### Identification of ligand binding sites in the 21 complete AlphaFold2 proteomes

We applied our ligand binding site search code to a total of 365,198 structures (representing 362,311 proteins) generated by AlphaFold2 [9], downloaded on 07/23/21 (https://alphafold.ebi.ac.uk/download).

All structures were treated identically, with the single exception of a “hypothetical repeat protein” from *Leishmania infantum* (UniProt accession: <u>**A4HXX6**</u>) which contains an extraordinary number of 6-histidine repeats arrayed in alpha helices in the AlphaFold2 structure. Running the code with our default cutoff of 8 Å for clustering into regions yields a single gigantic region so large that the code did not complete even after a week of run-time; for this particular case, therefore, the cutoff was changed to 6 Å. No binding sites were found. To process all of the other AlphaFold2 structures required a total of ^~^1200 CPU hours and so was run using locally accessible high performance computing resources.

For presenting results with a logical ordering of the 21 organisms, we generated a phylogenetic tree using the NCBI Taxonomy Browser (https://www.ncbi.nlm.nih.gov/Taxonomy/CommonTree/https://www.cmt.cgi).

### Comparing identified binding sites with annotated ligand binding sites in UniProt

To compare our identified ligand binding sites with known binding sites, the reference proteomes for the 21 organisms studied here were downloaded from the online protein database UniProt [43] as tab-separated files (accessed 9/3/21). These data files were then parsed to extract relevant information with a custom R script. Since information can sometimes be inconsistently annotated in UniProt entries, we attempted to capture the largest amount of data possible by designing the script to extract information from each of the following sections: “Cofactor”, “Sites”, “Region”, “Domain”, and “Zinc finger”. The most important of these sections is the “Sites” section since it explicitly identifies a variety of ligand and/or metal binding sites and provides lists of the exact coordinating residue numbers; in the present study, we searched for the following ligands: 4Fe-4S, 3Fe-4S, 2Fe-2S, Fe and Zn, with the latter two representing single metal binding sites; in unusual cases where instead of Fe or Zn it was listed as divalent metal cation, we searched the “Cofactor” section to determine that it was a Fe or Zn ions. To get a more complete assessment of Zn binding sites, we additionally examined the “Region”, “Domain” and “Zinc finger” sections, searching in the former two for the key words “Zinc” or “C*H*-type”, where the “*” ensures that we capture all possible combinations of cysteine and histidine coordination in zinc fingers. These latter sections provide additional sites but provide less specific information than is found the “Sites” section: rather than explicitly identifying the exact coordinating residues, they give a range of residues (e.g. “residues 760-800”) that contain the annotated binding site.

Recall rates were calculated for Fe-S clusters in the following way. We compared the ligand and coordinating residues identified by our method with those explicitly listed in the “Sites” section and required that all be matched exactly. Recall rates were for calculated for Zn-binding sites in the same way, but in cases where the “Sites” section was absent, we required that all coordinating residues identified by our method be contained within the range of residues entered in the “Region”, “Domain” and “Zinc finger” sections. We note that in order to properly assess our method’s ability to recall annotated binding sites, we exclude from our analysis any binding sites annotated in UniProt that involve coordinating residues that are not explicitly listed in our 12 ligand types. Having compared all identified binding sites for a given organism with all binding sites annotated within UniProt, we calculate the recall rate (also known as the sensitivity), R, as TP / (TP + FN) where TP and FN are the numbers of true positives and false negatives, respectively.

### Determining whether a predicted ligand binding site involves a region with no structural homolog

In an attempt to identify truly novel ligand binding sites built by AlphaFold2, we used tools provided in HH-suite 3.1.0 [30] to eliminate all binding sites for which homologous structures could be found in the RCSB. To do this, “hhblits” was first used to generate a multiple sequence alignment by searching against the Uniclust30 database [44]. We then ran “hhsearch” against the PDB70 database [30] using the multiple sequence alignment “.a3m” output from “hhblits”. This search for homologous structures was made as “greedy” as possible by adding the flag “ `-Z 10000`” to allow as many as 10,000 homologs to be listed in the output. We then wrote a script to eliminate all identified binding sites for which any of the coordinating residues were contained within the alignment produced by any homologous structure with an E-value of 1e-5 or better. For both the Uniclust30 and PDB70 databases, we took care to use identical versions used in the development of AlphaFold2 [8]. For Uniclust30, this is the August 2018 release (https://www.user.gwdg.de/~compbiol/uniclust/2018_08/); for PDB70, this was the May 13, 2020 release.

### Comparing identified binding sites with experiment and other predictions in the literature

For comparison with previous studies aiming to identify Fe-S containing proteins in E. coli, the predicted lists of proteins identified by Estellon et al. [21] were downloaded from the IronSulfurProteHome webserver (http://biodev.extra.cea.fr/isph/db/ECOLI.Fe-S.csv) and those identified by Valasatava *et al.* [22] were downloaded from the supporting information of the original publication. Since we compare only the proteins and not the predicting coordinating residues, we created non-redundant lists of proteins from each of these methods and our list of *E. coli* proteins identified as containing an Fe-S cluster, and then calculated the intersections between each of these three lists. We graphically represent the associations between these three Fe-S cluster proteomes as a Venn diagram using Microsoft PowerPoint.

For the comparisons with chemoproteomic data, lists of chemically reactive cysteine residues were downloaded for three human cancer cell proteomes from the same study ([32]; HCT116 cells from Supplementary Table 6, HEK293T cells from Supplementary Table 7 and PaTu-8988T cells from Supplementary Table 8) and for *E. coli* ([23], Supplementary Table 3). For each organism, we created a non-redundant list of all proteins that contained at least one chemically reactive cysteine and then determined the total number of cysteines contained in these proteins. We then constructed three subsets from the cysteines found within these proteins: (1) all “highly reactive” cysteines in the chemoproteomics data listed above, (2) cysteines identified here as coordinating residues in ligand binding sites, and (3) cysteines annotated in UniProt as coordinating residues in ligand binding sites. We then sought to assess the extent of overlap between these three subsets using one-tailed Fisher’s Exact tests. In comparing the cysteines identified here with the highly reactive cysteine subset, we tested the hypothesis that the two subsets segregate from each other more than expected by chance; in comparing the cysteines identified here with the UniProt subset, we tested instead the hypothesis that the two subsets overlap more than expected by chance. All tests were performed using the “fisher.test” function in the statistical programming language R [45]. We then created Euler (i.e. area-proportional Venn) diagrams illustrating the extent to which these three subsets of residues overlapped with each other using the “eulerr” library [46].

### Figure generation

Bar charts, histograms, and scatterplots were made in the statistical programming language R with the following packages: “ggplot2” [47], “gridExtra” [48], “scales” [49], “plotly” [50] and “eulerr” [46]. Images of proteins were made in VMD [51].

## Acknowledgments

The authors thank John Tworek and Venkata Sanaboyana for their help in the early stages of this project. This research was supported by a NIH grant (R35 GM122466) to AHE; ZJW was supported in part by a grant from Integrated DNA Technologies, Coralville, IA. This research was also supported in part through computational resources provided by The University of Iowa.

## Author Contributions

Zachary Wehrspan: Software, Formal analysis, Investigation, Data Curation, Writing – Original Draft, Visualization, Funding acquisition. Robert McDonnell: Software, Formal analysis, Investigation, Data Curation, Writing – Original Draft, Visualization. Adrian H. Elcock: Conceptualization, Methodology, Software, Investigation, Writing = Original Draft, Supervision, Project Administration, Funding acquisition.

## Data availability

A full listing of all ligand binding sites identified here can be found in Supplementary File 1. Full coordinate files in PDB format for all placed ligands can be found in Supplementary File 2.

