## Supplemental Figures for "Identification of Iron-Sulfur (Fe-S) and Zn-binding Sites Within Proteomes Predicted by DeepMind’s AlphaFold2 Program Dramatically Expands the Metalloproteome"

### Supplementary Figure Legends

Figure S1 Distribution of the RMSDs of the fits to the AlphaFold2 structures for all twelve ligand types that were successfully placed. Only cases in which all coordinating sidechains had pLDDT values of greater than 70 are recorded. The black, dashed line on each plot demarks the 0.5 Å RMSD threshold used to determine acceptable fits.

Figure S2 Fractional abundance of (a) identified ligand-binding sites and (b) identified ligand-binding proteins for all organisms. Only cases in which all coordinating sidechains had pLDDT values of greater than 70 are recorded

Figure S3 Examples of 4Fe-4S cluster binding sites identified with three coordinating cysteine residues. Protein chains are shown as cartoon colored using AlphaFold2's pLDDT score on a spectrum of red (low confidence) to blue (high confidence), 4Fe-4S clusters are shown as spheres with iron atoms colored pink and sulfur atoms colored yellow, and coordinating residues are shown as licorice colored by element with coordinating bonds labeled. (a) a binding site with a possible fourth coordinating cysteine residue (red arrow) that is too far away to be identified as part of the binding site using a 0.5 Å RMSD threshold. (b) a binding site with the fourth coordinating position exposed to bulk solvent. This image was prepared using VMD [51].

Figure S4 Examples of Zn-binding sites identified with three coordinating histidine residues. Visualization scheme is the same as used in Figure S3. (a) a binding site with a nearby fourth histidine residue that is not quite in a coordinating geometry. (b) a binding site with a nearby aspartate residue as a candidate fourth coordinating residue. (c) a binding site with the fourth coordinating position exposed to bulk solvent.

Figure S5 Venn diagram showing the extent of overlap between Fe-S cluster-binding proteins identified here in *E. coli* with those identified in previous prediction studies. Those identified here ("This Work") are represented by the red circle, those predicted by Estellon et al. [21] by the green circle, and those predicted by Valastava et al. [22] by the blue circle.

Figure S6 Distribution of the RMSDs of the fits to the AlphaFold2 structures for all twelve ligand types that were successfully placed for: (a) binding sites already annotated in UniProt, and (b) binding sites not annotated in UniProt. The blank panels in (a) are the result of two factors: (1) the rarity with which some ligand types are documented in UniProt (e.g. "2Fe-2S $Cys_3$ $Asp_1$" and "Zn $His_4$"), and (2) priority being given by our scheme to a different ligand type, e.g. UniProt-annotated "3Fe-4S $Cys_3$" binding sites are identified instead as "4Fe-4S $Cys_3$" binding sites).

Figure S7 Distribution of the RMSDs of the fits to the AlphaFold2 structures for all twelve ligand types that were successfully placed for: (a) binding sites already annotated in UniProt, or for which a structural homolog is identified with hhsearch, and (b) binding sites not annotated in UniProt and for which no structural homolog is identified with hhsearch. The blank panels that appear in (b) are the result of the strict criteria of using UniProt plus hhsearch to filter what we consider novel identified ligand binding sites.

Figure S8 Comparison of cysteines in the human proteome that are: (a) chemically reactive in experiment, (b) identified here in ligand binding sites, and/or (c) annotated in UniProt. All comparisons are shown as Euler diagrams with the outer circle representing the total number of cysteines in proteins that contained at least one reactive cysteine in the isoTOP-ABPP experiments [32], blue circles represent the subset of cysteines that are classed as "highly reactive" in the -ABPP experiments ("Kuljanin *et al*. Union"), red circles represent the subset of cysteines that are identified here as being parts of ligand binding sites, and grey circles represent the subset of cysteines that are annotated in UniProt as members of ligand binding sites ("UniProt"). (a) shows results obtained when all binding sites identified here are included (i.e. no pLDDT cutoff, (b) shows results obtained when only binding sites identified here for which all coordinating residues have pLDDT scores > 90 are included, (c) shows results obtained when only binding sites identified here for which all coordinating residues have pLDDT scores > 70 are included, and when all chemically reactive cysteines are included.

Figure S9 Comparison of cysteines in the *E. coli* proteome that are: (a) chemically reactive in experiment, (b) identified here in ligand binding sites, and/or (c) annotated in UniProt. All comparisons are shown as Euler diagrams with the outer circle representing the total number of cysteines in proteins that contained at least one reactive cysteine in the isoTOP-ABPP experiments [23], blue circles represent the subset of cysteines that are classed as “highly reactive” in the experiments (“Wang *et al*.”), red circles represent the subset of cysteines that are identified here as being parts of ligand binding sites, and grey circles represent the subset of cysteines that are annotated in UniProt as members of ligand binding sites (“UniProt”). (a) shows results obtained when all binding sites identified here are included (i.e. no pLDDT cutoff, (b) shows results obtained when only binding sites identified here for which all coordinating residues have pLDDT scores > 90 are included, (c) shows results obtained when only binding sites identified here for which all coordinating residues have pLDDT scores > 70 are included, and when all chemically reactive cysteines are included.

Figure S10 A case highlighting AlphaFold2’s rare indecision when building metal binding sites in overlapping fragments. Visualization scheme is the same as used in Figure S3. (a) fragment O43149_F3 contains a single zinc coordinated by 4 cysteine residues (“Zn $Cys_4$”). (b) fragment O43149_F4 (and 5 subsequent fragments) contain a single zinc coordinated by 2 cysteine and two histidine residues (“Zn $Cys_2$ $His_2$”).

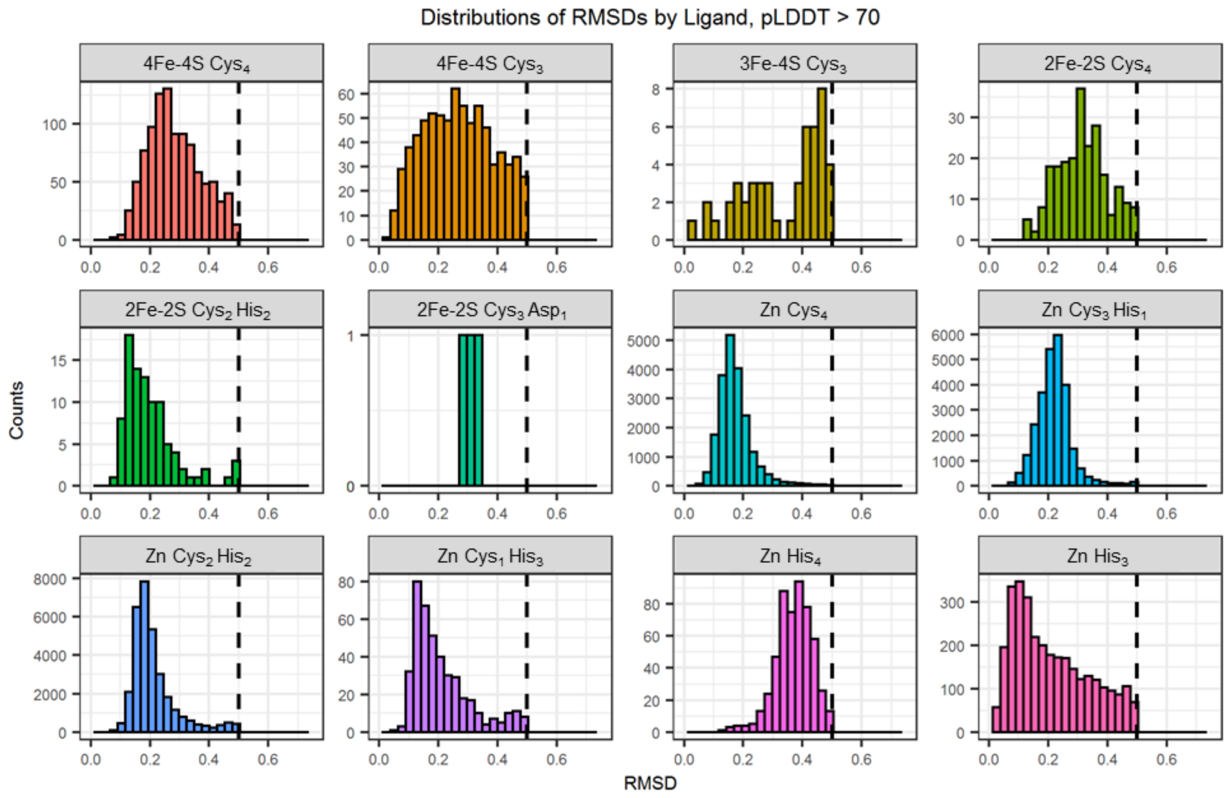

Distributions of RMSDs by Ligand, pLDDT > 70
4Fe-4S Cys4
4Fe-4S Cys3
3Fe-4S Cys3
2Fe-2S Cys4
2Fe-2S Cys2 His2
2Fe-2S Cys3 Asp1
Zn Cys4
Zn Cys3 His1
Zn Cys2 His2
Zn Cys1 His3
Zn His4
Zn His3
Counts
RMSD


**Figure S1**

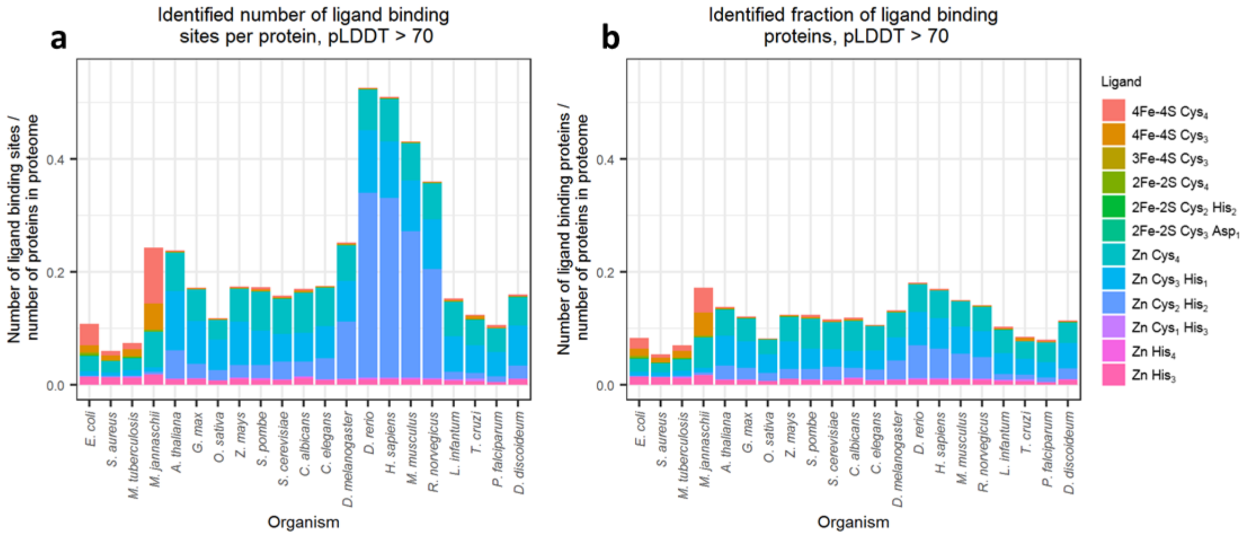

a
Identified number of ligand binding sites per protein, pLDDT > 70
Number of ligand binding sites / number of proteins in proteome
0.0
0.2
0.4
E. coli
S. aureus
M. tuberculosis
M. jannaschii
A. thaliana
G. max
O. sativa
Z. mays
S. pombe
S. cerevisiae
C. albicans
C. elegans
D. melanogaster
D. rerio
H. sapiens
M. musculus
R. norvegicus
L. infantum
T. cruzi
P. falciparum
D. discoideum
Organism
b
Identified fraction of ligand binding proteins, pLDDT > 70
Number of ligand binding proteins / number of proteins in proteome
Organism
Ligand
4Fe-4S Cys4
4Fe-4S Cys3
3Fe-4S Cys3
2Fe-2S Cys4
2Fe-2S Cys2 His2
2Fe-2S Cys3 Asp1
Zn Cys4
Zn Cys3 His1
Zn Cys2 His2
Zn Cys1 His3
Zn His4
Zn His3


**Figure S2**

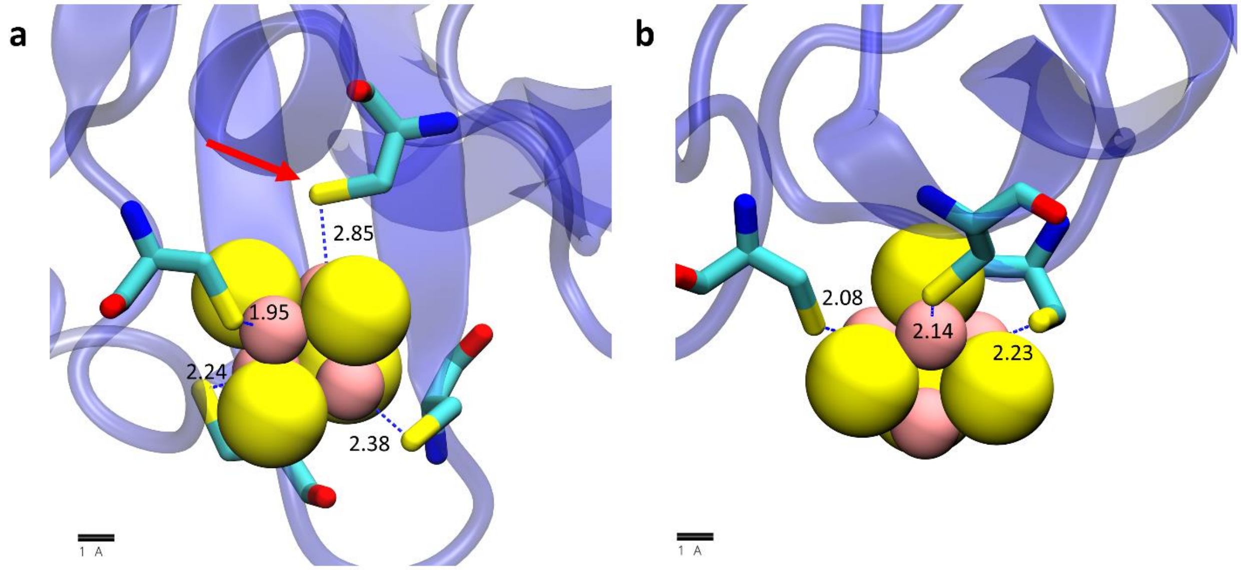


**Figure S3**

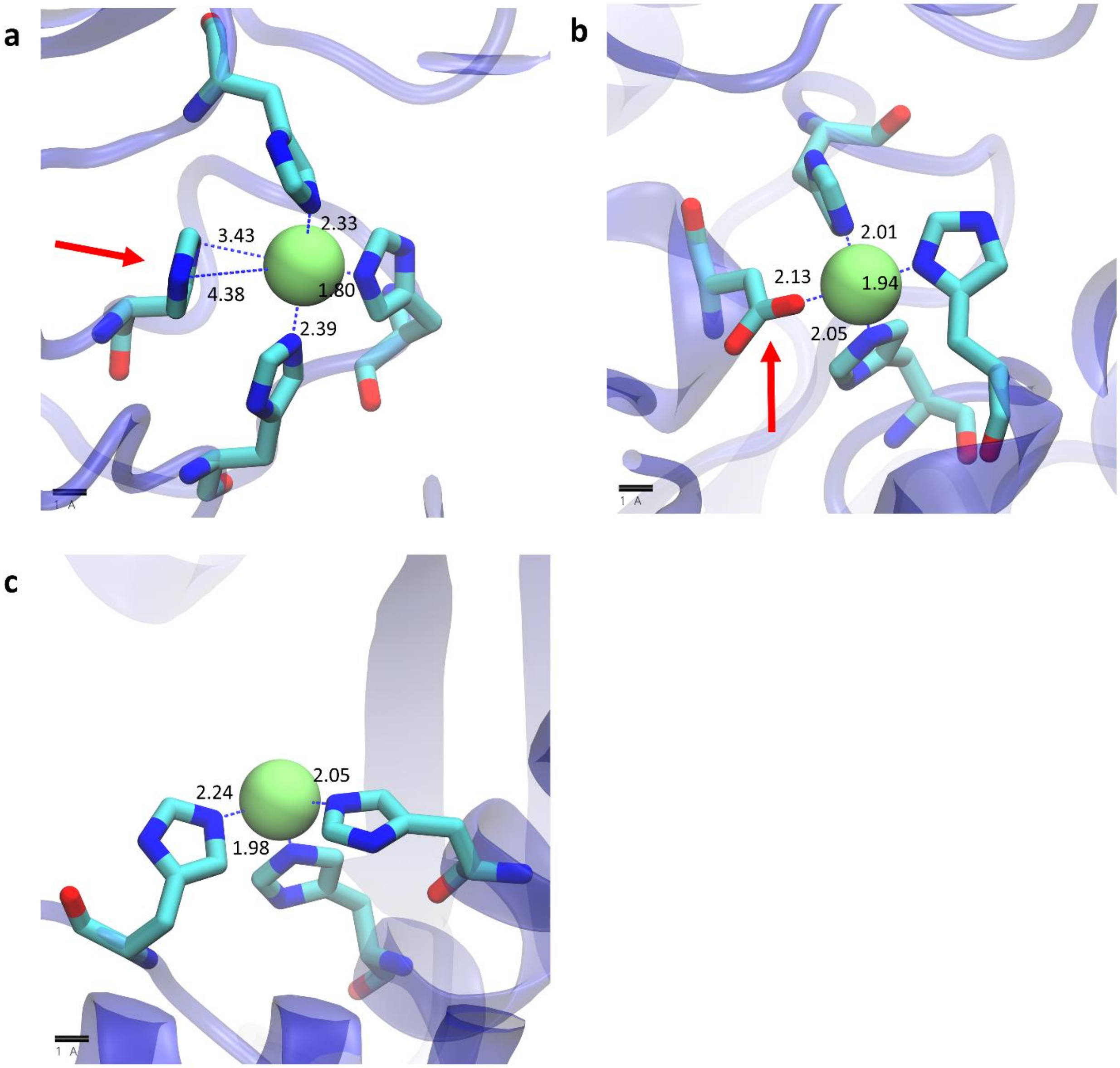


**Figure S4**

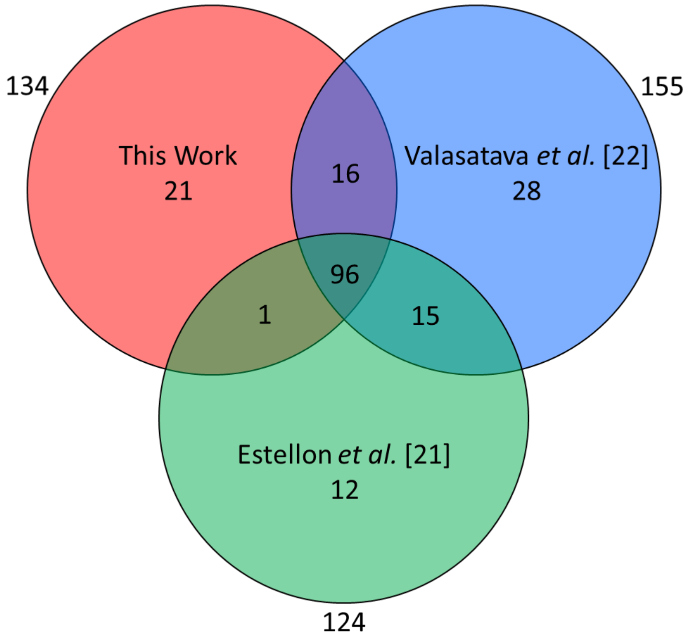
134
155
This Work
21
16
Valasatava *et al.* [22]
28
96
1
15
Estellon *et al.* [21]
12
124

**Figure S5**

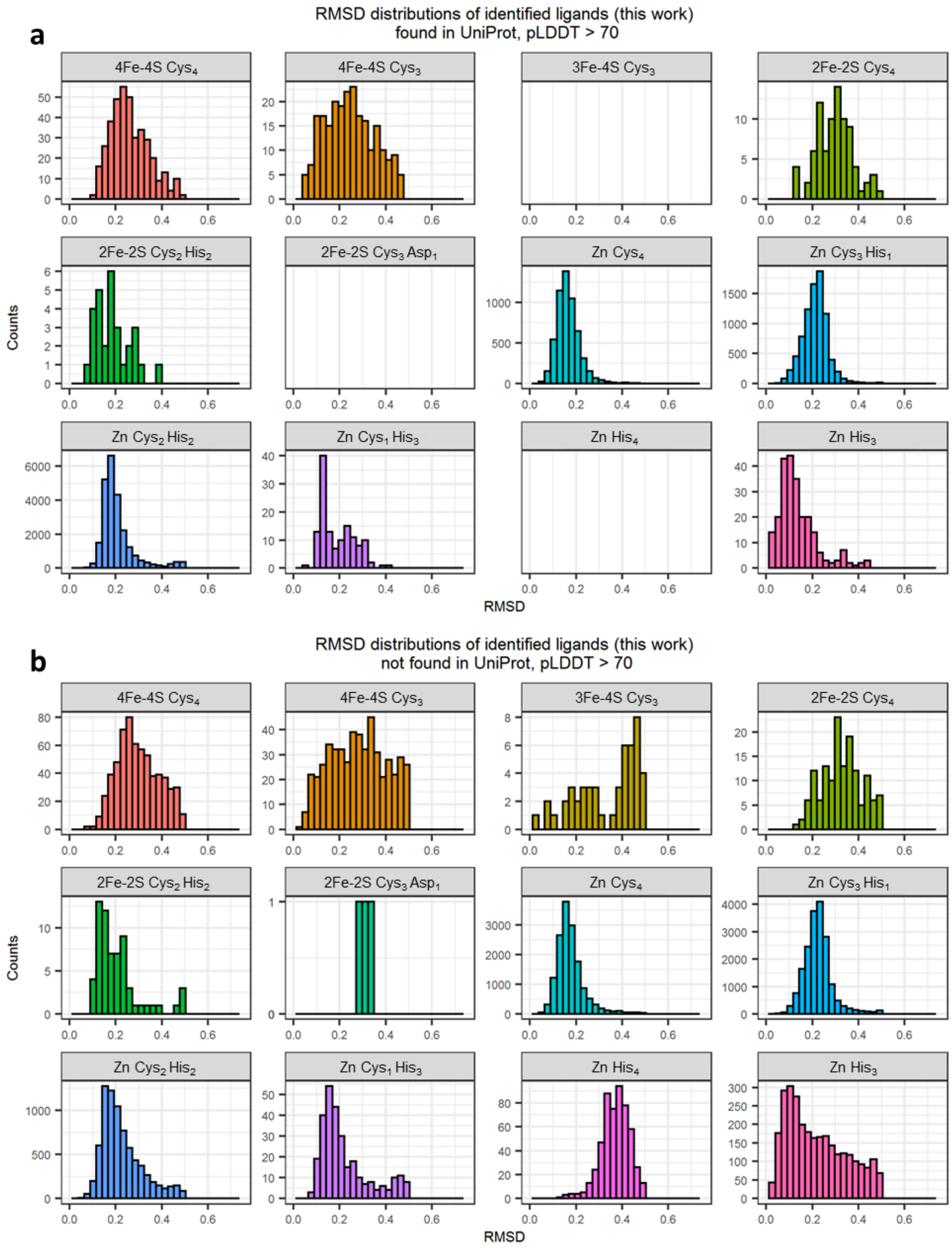
a
RMSD distributions of identified ligands (this work)
found in UniProt, pLDDT > 70
4Fe-4S $Cys_4$
4Fe-4S $Cys_3$
3Fe-4S $Cys_3$
2Fe-2S $Cys_4$
2Fe-2S $Cys_2$ $His_2$
2Fe-2S $Cys_3$ $Asp_1$
Zn $Cys_4$
Zn $Cys_3$ $His_1$
Zn $Cys_2$ $His_2$
Zn $Cys_1$ $His_3$
Zn $His_4$
Zn $His_3$
Counts
RMSD
b
RMSD distributions of identified ligands (this work)
not found in UniProt, pLDDT > 70
4Fe-4S $Cys_4$
4Fe-4S $Cys_3$
3Fe-4S $Cys_3$
2Fe-2S $Cys_4$
2Fe-2S $Cys_2$ $His_2$
2Fe-2S $Cys_3$ $Asp_1$
Zn $Cys_4$
Zn $Cys_3$ $His_1$
Zn $Cys_2$ $His_2$
Zn $Cys_1$ $His_3$
Zn $His_4$
Zn $His_3$
Counts
RMSD

**Figure S6**

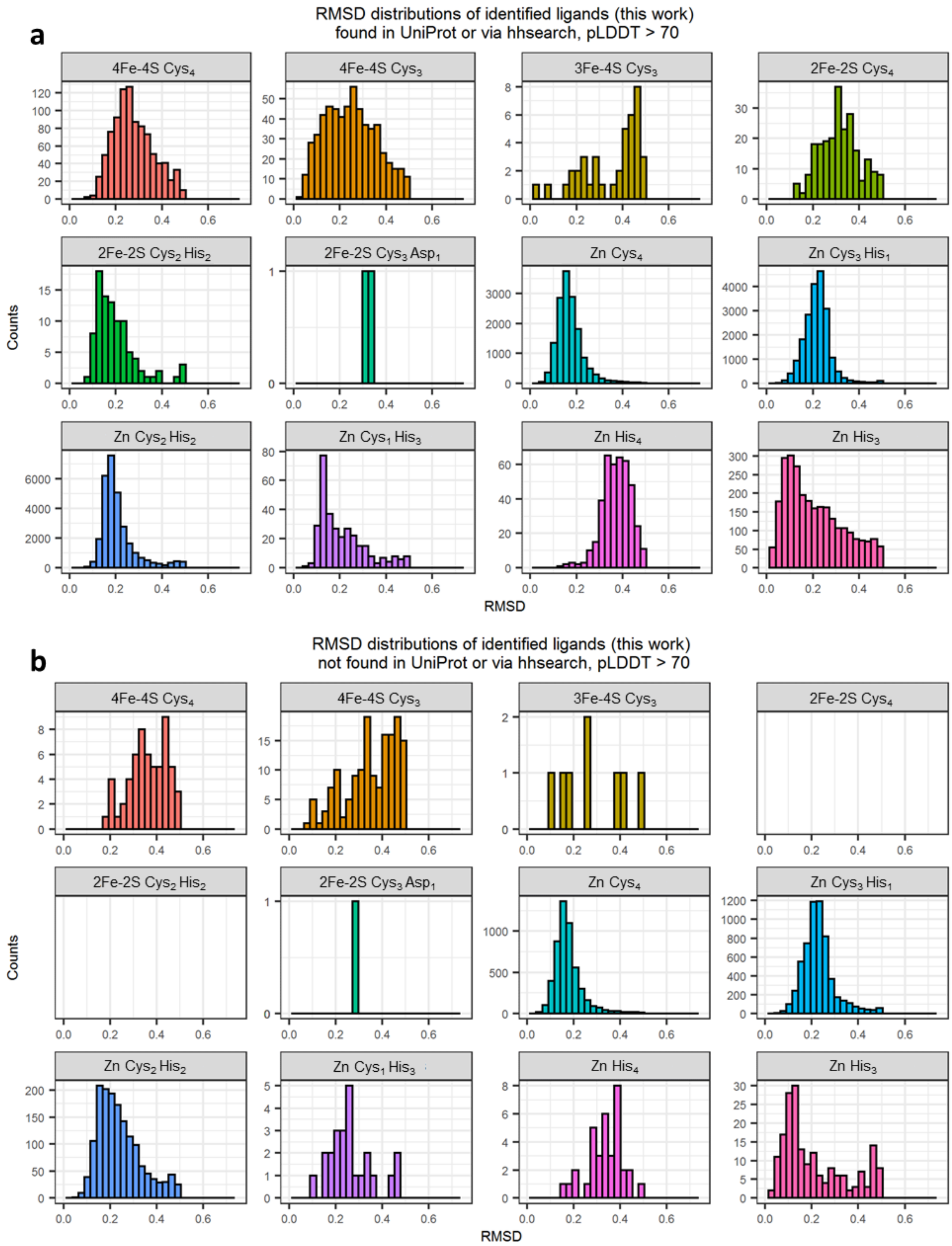
a
RMSD distributions of identified ligands (this work)
found in UniProt or via hhsearch, pLDDT > 70
4Fe-4S $Cys_4$
4Fe-4S $Cys_3$
3Fe-4S $Cys_3$
2Fe-2S $Cys_4$
2Fe-2S $Cys_2$ $His_2$
2Fe-2S $Cys_3$ $Asp_1$
Zn $Cys_4$
Zn $Cys_3$ $His_1$
Zn $Cys_2$ $His_2$
Zn $Cys_1$ $His_3$
Zn $His_4$
Zn $His_3$
Counts
RMSD
b
RMSD distributions of identified ligands (this work)
not found in UniProt or via hhsearch, pLDDT > 70
4Fe-4S $Cys_4$
4Fe-4S $Cys_3$
3Fe-4S $Cys_3$
2Fe-2S $Cys_4$
2Fe-2S $Cys_2$ $His_2$
2Fe-2S $Cys_3$ $Asp_1$
Zn $Cys_4$
Zn $Cys_3$ $His_1$
Zn $Cys_2$ $His_2$
Zn $Cys_1$ $His_3$
Zn $His_4$
Zn $His_3$
Counts
RMSD

**Figure S7**

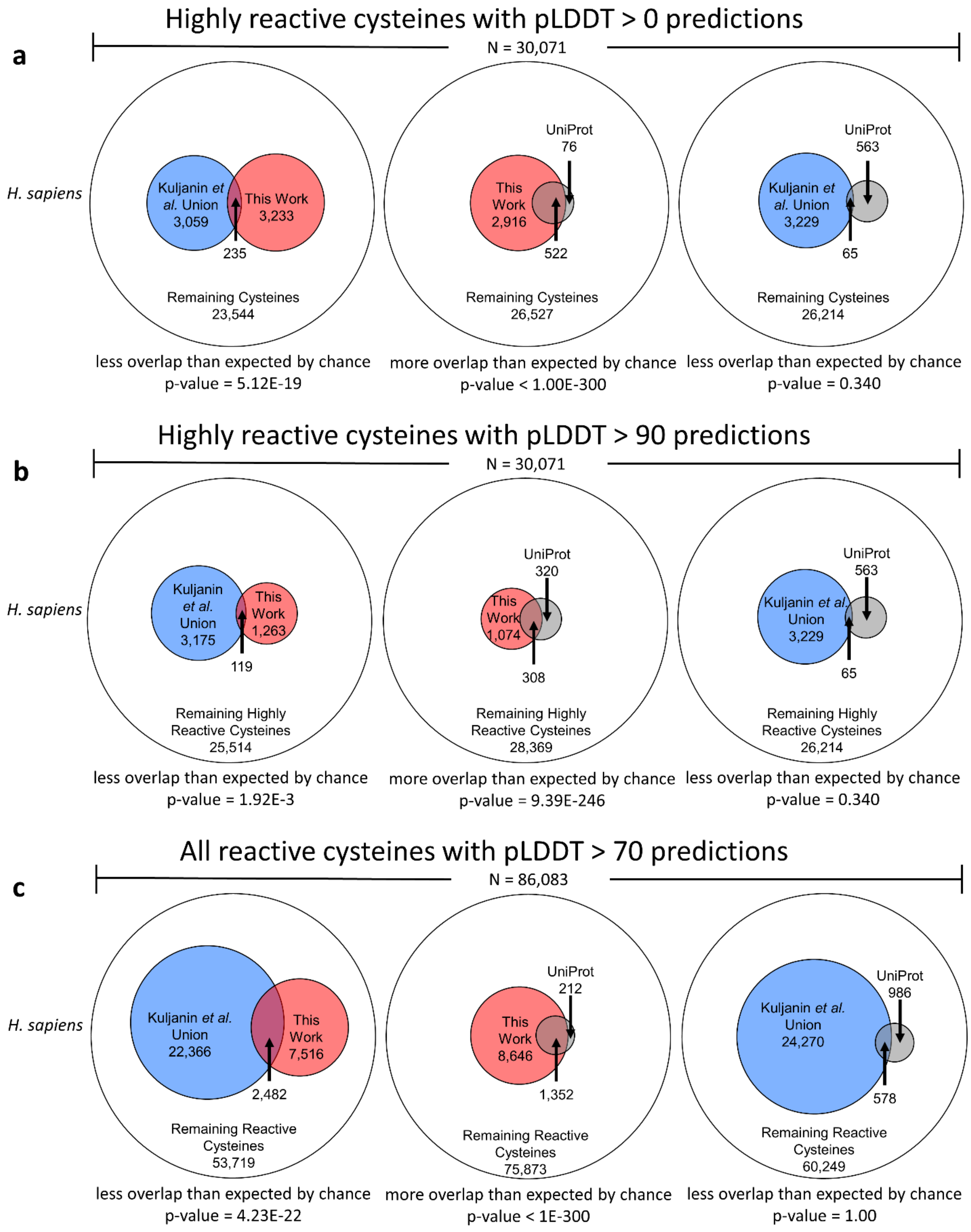

Highly reactive cysteines with pLDDT > 0 predictions
a
N = 30,071
H. sapiens
Kuljanin et al. Union 3,059
This Work 3,233
235
Remaining Cysteines 23,544
less overlap than expected by chance
p-value = 5.12E-19
UniProt 76
This Work 2,916
522
Remaining Cysteines 26,527
more overlap than expected by chance
p-value < 1.00E-300
UniProt 563
Kuljanin et al. Union 3,229
65
Remaining Cysteines 26,214
less overlap than expected by chance
p-value = 0.340
Highly reactive cysteines with pLDDT > 90 predictions
b
N = 30,071
H. sapiens
Kuljanin et al. Union 3,175
This Work 1,263
119
Remaining Highly Reactive Cysteines 25,514
less overlap than expected by chance
p-value = 1.92E-3
UniProt 320
This Work 1,074
308
Remaining Highly Reactive Cysteines 28,369
more overlap than expected by chance
p-value = 9.39E-246
UniProt 563
Kuljanin et al. Union 3,229
65
Remaining Highly Reactive Cysteines 26,214
less overlap than expected by chance
p-value = 0.340
All reactive cysteines with pLDDT > 70 predictions
c
N = 86,083
H. sapiens
Kuljanin et al. Union 22,366
This Work 7,516
2,482
Remaining Reactive Cysteines 53,719
less overlap than expected by chance
p-value = 4.23E-22
UniProt 212
This Work 8,646
1,352
Remaining Reactive Cysteines 75,873
more overlap than expected by chance
p-value < 1E-300
UniProt 986
Kuljanin et al. Union 24,270
578
Remaining Reactive Cysteines 60,249
less overlap than expected by chance
p-value = 1.00


**Figure S8**

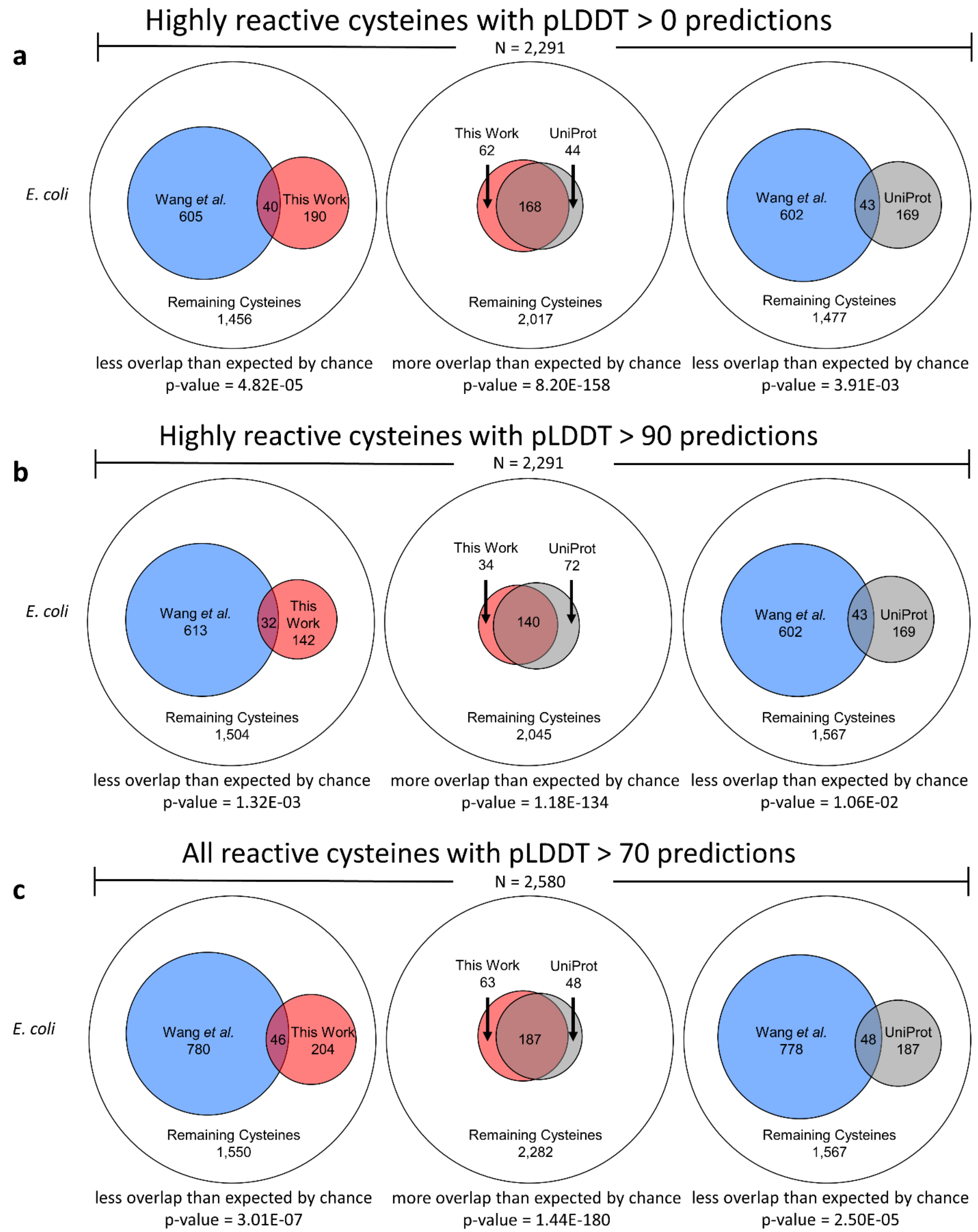
a
Highly reactive cysteines with pLDDT > 0 predictions
N = 2,291
E. coli
Wang et al. 605
40
This Work 190
Remaining Cysteines 1,456
less overlap than expected by chance
p-value = 4.82E-05
This Work 62
UniProt 44
168
Remaining Cysteines 2,017
more overlap than expected by chance
p-value = 8.20E-158
Wang et al. 602
43
UniProt 169
Remaining Cysteines 1,477
less overlap than expected by chance
p-value = 3.91E-03
b
Highly reactive cysteines with pLDDT > 90 predictions
N = 2,291
E. coli
Wang et al. 613
32
This Work 142
Remaining Cysteines 1,504
less overlap than expected by chance
p-value = 1.32E-03
This Work 34
UniProt 72
140
Remaining Cysteines 2,045
more overlap than expected by chance
p-value = 1.18E-134
Wang et al. 602
43
UniProt 169
Remaining Cysteines 1,567
less overlap than expected by chance
p-value = 1.06E-02
c
All reactive cysteines with pLDDT > 70 predictions
N = 2,580
E. coli
Wang et al. 780
46
This Work 204
Remaining Cysteines 1,550
less overlap than expected by chance
p-value = 3.01E-07
This Work 63
UniProt 48
187
Remaining Cysteines 2,282
more overlap than expected by chance
p-value = 1.44E-180
Wang et al. 778
48
UniProt 187
Remaining Cysteines 1,567
less overlap than expected by chance
p-value = 2.50E-05

**Figure S9**

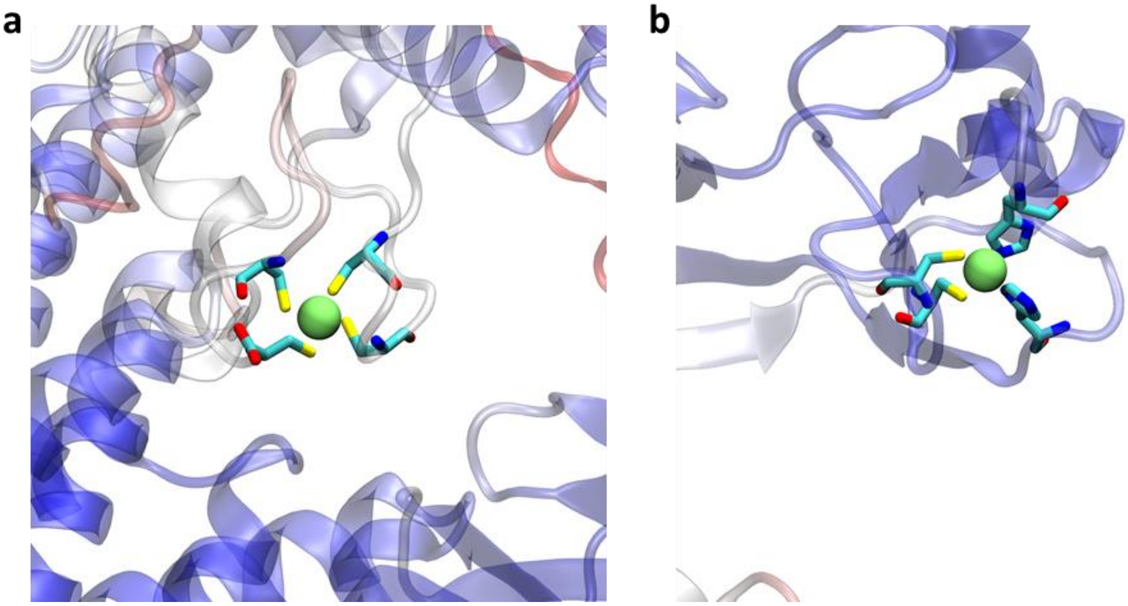


**Figure S10**
